# scIRES-seq reveals multimodal molecular heterogeneity in CTCs from small cell cervical cancer

**DOI:** 10.64898/2026.10.07.757279

**Authors:** Lea Kubitz, Katharina Bauer, Diana Ordoñez-Rueda, Maximilian Schönung, Valentin Maurer, Panna Lajer, Stephen J. Clark, Daniel Hübschmann, Maria-Veronica Teleanu, Wolf-Karsten Hofmann, Steffen Schmitt, Jan-Philipp Mallm, Stefan Fröhling, Tobias Boch, Daniel B. Lipka, Mark Hartmann

## Abstract

Circulating tumor cells (CTCs) can provide a window into cellular and molecular mechanisms of cancer dissemination. However, the rarity and heterogeneity of CTCs complicate reliable identification and molecular characterization. Here, we introduce scIRES-seq, a framework linking immunophenotype, genetic variation, epigenomic profiles and gene expression within individual cells. In this proof-of-concept study, we coupled scIRES-seq to real-time imaging spectral flow cytometry and applied it to CTCs from a patient with metastatic cervical small cell neuroendocrine carcinoma. Tumor-matched copy-number alterations and targeted somatic variants established CTC identity, supported by DNA methylation profiles concordant with metastatic tissue and transcriptional features consistent with neuroendocrine differentiation. Genetic and DNA methylation analyses identified two largely concordant CTC subpopulations that were not clearly distinguished by their transcriptomes, highlighting the complementary information gained through single-cell multi-omics profiling. These findings demonstrate the feasibility of integrated CTC identification and characterization and provide a framework for investigating relationships between genetic diversity and molecular states in rare, heterogeneous cell populations.

**Highlights:**

- scIRES-seq links single-cell indexing of cellular phenotypes, genetic variation, DNA methylation and gene expression within individual patient-derived CTCs.
- Tumor-matched genetic alterations, complementary DNA methylation and transcriptional signatures establish the tumor identity of rare circulating cells.
- CTCs harbor largely concordant genetic and epigenetic subpopulations that are not clearly distinguished by their transcriptomes, highlighting the complementary information gained through single-cell multi-omics profiling.

## INTRODUCTION

Circulating tumor cells (CTCs) are malignant cells released from primary or metastatic lesions into the bloodstream, where they can occur individually or in clusters. Their accessibility through blood sampling provides an opportunity to investigate cancer dissemination and the molecular composition of advanced disease. CTC enumeration has established prognostic value in entities like metastatic breast cancer, while transplantation studies have demonstrated that selected CTC populations can initiate metastases.^1–3^ These observations motivate molecular investigation of CTCs beyond enumeration, without implying that every circulating tumor cell is capable of establishing a metastatic lesion.

Characterizing CTCs is challenging because they are rare among blood cells and heterogeneous in both genetic composition and molecular phenotype. Studies have documented clonal diversity and variation in epithelial and mesenchymal expression programs, including changes in their representation during treatment and disease progression.^4,5^ Enrichment strategies can influence which populations are recovered and affect resulting molecular readouts.^6^ Moreover, CTCs may differ because they belong to genetically distinct populations, because cells with particular pre-existing molecular features are preferentially selected, or because individual cells change their molecular state. A key question is whether these molecular differences coincide with genetic subpopulations or occur among cells with similar measured genetic profiles. Reliable tumor-cell isolation and measurements linking genetic identity with molecular state in the same cells are therefore central to interpreting CTC heterogeneity.

DNA methylation provides an additional dimension for investigating CTC identity and heterogeneity. Early studies detected promoter methylation in enriched CTC fractions, followed by targeted analyses demonstrating methylation differences between individual CTCs.^7,8^ Genome-wide profiling subsequently identified methylation differences between single CTCs and CTC clusters from breast cancer, complemented by functional experiments linking cluster-associated methylation changes to metastatic seeding in models.^9^ Studies of lung-cancer CTC preparations further described tumor-associated methylation patterns and relationships with matched tumor tissue.^10,11^ These findings establish the relevance of CTC methylation profiling while highlighting the different information obtained from enriched fractions, pooled cells and individual cells.

Measurements obtained from the same cell allow genetic alterations, methylation patterns and transcriptional programs to be examined in direct correspondence. This approach has already been applied to CTCs: Chen and colleagues combined single-cell methylomes with inferred copy-number alterations across several cancer types, and Guo and colleagues obtained paired methylome and transcriptome measurements from individual prostate CTCs.^12,13^ Such measurements enable assessment of whether molecular differences coincide with genetically defined subpopulations. Linking them to recorded immunophenotypes and cell images additionally connects the properties used during isolation with subsequent molecular evidence of tumor identity.

Established single-cell methods provide complementary foundations for this integration. G&T-seq enables physical separation of genomic DNA and messenger RNA, while scM&T-seq connects methylome and transcriptome measurements within individual cells using single-cell bisulfite sequencing (scBS-seq) and Smart-seq2.^14–17^ MutaSeq complements transcriptome profiling through targeted amplification of expressed mutation-bearing loci, supporting genetic assignment of individual cells.^18^

Here, we introduce scIRES-seq, an integrated framework combining adapted versions of these methods with index sorting to connect immunophenotype, genetic information, DNA methylation and transcription within individual cells. We couple scIRES-seq to real-time imaging spectral flow cytometry (RTI-SFC) applying the image-enabled cell sorting technology^19^, using SFC for cell sorting and retaining the acquired images for subsequent characterization of sorted events. We apply this framework to CTCs from a patient in palliative care with metastatic cervical small cell neuroendocrine carcinoma, a rare and aggressive malignancy.^20^ Using molecular data from an earlier metastatic biopsy as a reference, we assess tumor identity through copy-number alterations, targeted somatic variants, methylation profiles and transcriptional features. We further identify largely concordant genomic and epigenomic CTC subpopulations that are not clearly separated by the available transcriptome analysis. This proof-of-concept study illustrates how linked measurements can support CTC identification and reveal complementary aspects of molecular heterogeneity within a clinical sample.

## RESULTS

In this proof-of-concept study, we used scIRES-seq to investigate CTC identity and molecular heterogeneity in a patient with advanced cervical small cell neuroendocrine carcinoma. We selected this patient based on the extensive metastatic disease, anticipating recovery of sufficient CTCs for single-cell multi-omics analysis. Published single-cell data have demonstrated transcriptional heterogeneity in cervical small cell carcinoma, providing a rationale for investigating its relationship to genetic and epigenetic variation.^21^ This study adds molecular information about a rare, aggressive malignancy for which research material is difficult to obtain.^20^

### Clinical course and sampling of metastatic cervical small cell neuroendocrine carcinoma

The patient was diagnosed with FIGO stage IV poorly differentiated small cell neuroendocrine carcinoma of the cervix (G3; Ki-67, 95%) with bilateral adrenal metastases. Despite transient responses to multiple systemic treatments, including platinum–etoposide chemotherapy, immunotherapy and an investigational DLL3×CD3 bispecific antibody, disease progressed to involve the brain, leptomeninges, orbits and liver. Pelvic and intracranial radiotherapy and palliative orbital irradiation were administered during the disease course. Following further progression, care transitioned to supportive palliative management, and the patient died from progressive disease.

The patient had been enrolled in the NCT-MASTER precision oncology program.^22^ Whole-genome sequencing (WGS) of an earlier subcutaneous arm metastasis and blood as a germline reference, together with EPIC methylation array data from the same metastasis, provided molecular reference profiles (all 4 months before death). A further blood sample was collected for CTC analysis three weeks before death, during advanced terminal disease (Fig. 1A).

**Figure 1.**
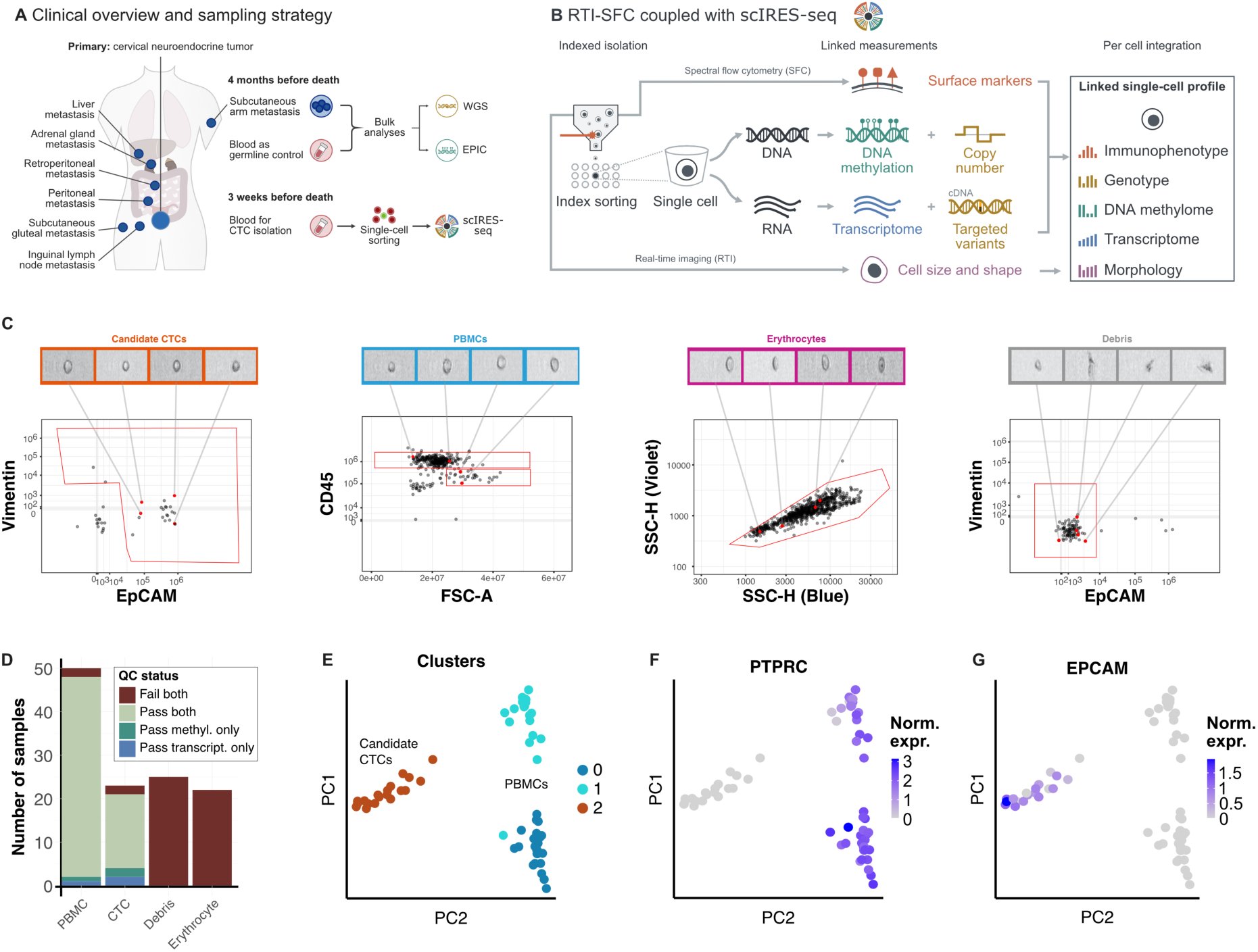
Isolation of circulating tumor cells and multiomics single-cell profiling with scIRES-seq. **A** Clinical overview and sampling strategy for the index patient. Anatomical schematic showing the distribution of metastatic lesions, the biopsy sampling site, sample collection timepoints relative to death and obtained data modalities. WGS, whole genome sequencing; scIRES-seq, single-cell immunophenotype/RNA/epigenome/SNV-sequencing. **B** Overview of scIRES-seq workflow. Viable single cells were sorted onto a 384-well plate with a real-time imaging spectral flow cytometer (BD FACSDiscover S8). Following cell lysis, DNA and mRNA were separated onto two plates with poly-dT magnetic beads. Genome-wide DNA methylation was assessed through scBS-seq. Single-cell full-length transcriptome libraries were obtained with an adapted SmartSeq2 protocol. Targeted primers were spiked in during cDNA amplification to increase sequencing coverage for genotyping at loci of interest. All molecular measurements of each cell were linked by indexing through the well coordinate on the 384-well plate. RTI, real-time imaging; SFC, spectral flow cytometry. **C** Gating strategy and representative images for sorted cell populations. Red polygons enclose the target population and were set based on marker expression (CD45, EpCAM, Vimentin) and scatter properties. Candidate CTCs and debris were sorted from the CD45-depleted fraction after MACS. PBMCs were sorted from the CD45-enriched fraction after MACS, with distinction of a CD45-high and CD45-intermediate population. Erythrocytes were sorted from unstained whole blood. Callouts show representative images from the Lightloss channel obtained from the FACSDiscover S8. Widths of the images are consistent between all events. The CellView technology does not provide scale bar information. For visualization, .fcs files were randomly downsampled to 1,000 raw events including the sorted events. EPCAM, epithelial cell adhesion molecule; FSC-A, forward scatter-area; SSC-H, side scatter-height; PBMC, peripheral blood mononuclear cell; MACS, magnetic-activated cell sorting. **D** Number of cells passing library quality control for methylome and transcriptome. Methylome libraries passed quality control with minimum 50% uniquely aligned reads, minimum 100,000 CpG sites covered and maximum 5% cytosine methylation in a non-CpG context. Transcriptome libraries passed quality control with minimum 10,000 raw reads, minimum 1,000 detected genes and maximum 10% mitochondrial gene counts. PBMCs comprised 48 single cells, an empty well as negative control and a 25-cell minibulk. Two candidate CTCs passed all metrics except for slightly too high CHH methylation (5.2% and 5.7%). Two further candidate CTCs passed all metrics except for intermediately elevated mitochondrial gene counts (13% and 15%). These cells were included or excluded in particular analyses based on the specific requirements. **E - G** Principal Component Analysis of single-cell transcriptomes of PBMCs and candidate CTCs. Color indicates Louvain cluster identity (**E**) or log-normalized gene expression of (**F**) PTPRC (encoding CD45) and (**G**) EPCAM.

### scIRES-seq links cell imaging and immunophenotype to matched methylome and transcriptome profiles

We isolated live candidate CTCs using real-time imaging spectral flow cytometry (RTI-SFC), linking cell images and immunophenotypes to subsequent molecular measurements through index sorting (Fig. 1B). Because epithelial and mesenchymal marker expression can vary among CTCs, we initially sought to recover candidates without positive selection for a tumor-associated marker.^23^ Exclusion of CD45-positive immune cells and CD235a-positive erythrocytes alone, however, yielded predominantly amorphous debris-like events in the target gate, without recognizable intact-cell morphology (Fig. 1C, “Debris”). Similar observations were made in additional samples from different tumor entities and disease stages (data not shown). Adding positive selection for EpCAM and/or vimentin yielded events with intact-cell morphology (Fig. 1C, “Candidate CTCs”).

We sorted 23 candidate CTCs directly into lysis buffer. As blood-cell references, we sorted 48 individual CD45-positive events from CD45-high and CD45-intermediate gates as well as a 25-cell PBMC minibulk and an empty well as technical controls. We additionally collected 25 debris-like events and 22 erythrocyte events (Fig. 1C). Imaging confirmed singlet morphology for all 23 candidate CTCs and 46 of the 48 individually sorted peripheral blood mononuclear cell (PBMC) events (Fig. S6). Index sorting linked each event’s images and fluorescence measurements to its well coordinate on a 384-well plate.

We generated matched single-cell transcriptome and bisulfite-based DNA methylome libraries using scIRES-seq, as described in the Methods. Shared well coordinates linked the molecular measurements to each other and to the corresponding imaging and immunophenotypic data, without requiring computational matching between separately profiled cells.

Both methylome and transcriptome profiles passed quality control for 46 PBMCs and 17 candidate CTCs (Fig. 1D). An additional two PBMCs and four candidate CTCs passed QC for a single modality—one PBMC and two candidate CTCs for each assay—and were retained for analyses of that modality. Two candidate CTCs failed both assays; both were EpCAM-negative and vimentin-positive. Neither the empty-well control nor any debris-like event passed methylome or transcriptome QC.

Transcriptome clustering separated the analyzed cells into three groups (Fig. 1E). Expression of PTPRC, encoding CD45, and EPCAM was consistent with the PBMC and candidate-CTC sorting populations, respectively (Fig. 1F–G). Together, these results demonstrate recovery of intact candidate CTCs and generation of matched methylome and transcriptome profiles for most sorted candidates.

### Complementary molecular measurements confirm the tumor origin of candidate CTCs

We next examined whether the recovered candidates carried molecular features of the patient’s tumor. Copy-number variation (CNV) profiles inferred from the single-cell methylome data were compared to WGS of a metastatic lesion. The candidate-CTC consensus, calculated as the median normalized read ratio per genomic bin, closely resembled the metastatic profile across the genome (Fig. 2A, left). Both profiles contained numerous corresponding copy-number gains and losses. In contrast, the PBMC consensus and bulk blood reference showed no major copy-number alterations (Fig. 2A, right).

**Figure 2.**
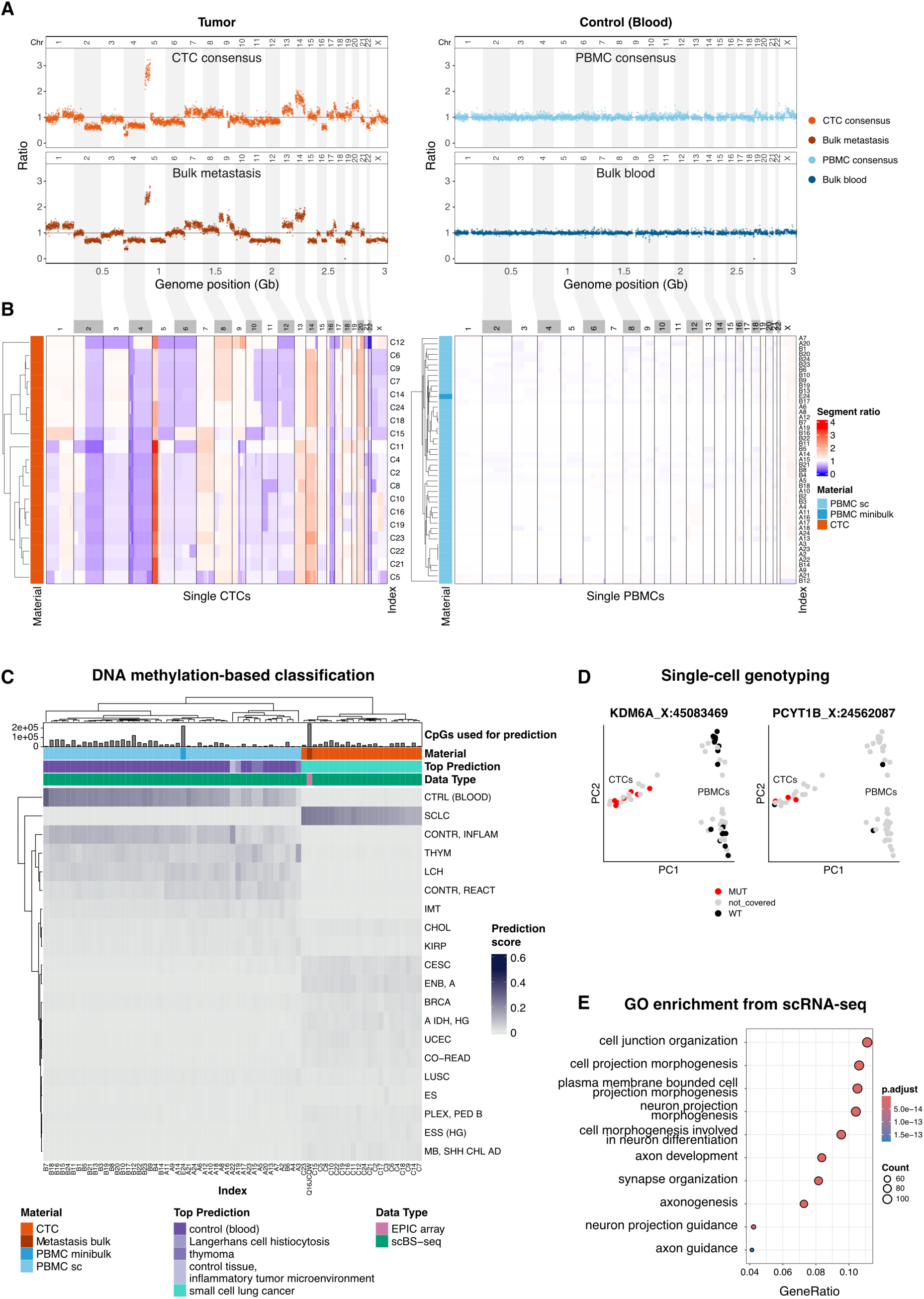
Multimodal confirmation of tumor origin for candidate CTCs. **A** Copy-number analysis based on bulk WGS data (blood and metastasis) and scBS-seq data (candidate CTCs, PBMCs). Color indicates sample type. Points represent the read count per genomic bin normalized to the average bin read count of each sample, using the Variable Binning algorithm and GC bias correction (copyKit package, 500 kb resolution). For PBMCs and candidate CTCs, the consensus was calculated as the median ratio per bin across single cells. Ratios below 1 indicate losses; ratios above 1 indicate gains. White and grey rectangles indicate chromosomes. **B** Single-cell copy-number profiles for single cells summarized in A. Bin ratios were segmented using the Circular Binary Segmentation algorithm on the combined CTC and PBMC dataset. Gains (ratio > 1) are shown in red; losses (ratio < 1) in blue. Rows were hierarchically clustered using Euclidean distance and complete linkage. Row annotations indicate index and sample material. **C** Methylation-based entity prediction using the pan-cancer classifier of the crossNN framework. Heatmap color indicates the prediction score (range: 0-1). Of 178 methylation classes, only those ranking in the top 5 for at least one sample are displayed. Input data were scBS-seq (PBMCs and CTCs) and EPIC array (metastasis). Column annotations indicate sample material, top-predicted methylation class, data type, and number of CpGs overlapping classifier input features. Columns and rows were hierarchically clustered using Euclidean distance and complete linkage. CTRL (BLOOD), control (blood); SCLC, small cell lung cancer; CONTR INFLAM, control tissue, inflammatory tumor microenvironment; THYM, thymoma; LCH, Langerhans cell histiocytosis; CONTR REACT, control tissue, reactive tumor microenvironment; IMT, inflammatory myofibroblastic tumor; CHOL, cholangiocarcinoma; KIRP, renal papillary cell carcinoma; CESC, cervical squamous cell carcinoma and endocervical adenocarcinoma; ENB A, esthesioneuroblastoma subclass A; BRCA, breast invasive carcinoma; A IDH HG, IDH glioma subclass high-grade astrocytoma; UCEC, uterine corpus endometrial carcinoma; CO-READ, colorectal adenocarcinoma; LUSC, lung squamous cell carcinoma; ES, epithelioid sarcoma; PLEX PED B, plexus tumor subclass paediatric B; ESS (HG), endometrial stromal sarcoma (high grade); MB SHH CHL AD, medulloblastoma subclass SHH A (children and adults). **D** Principal component analysis of single-cell transcriptomes. Color indicates mutation status at tumor-specific mutation loci (KDM6A chrX:45083469 and PCYT1B chrX:24562087). A cell was classified as mutated if at least one read carried the tumor-specific SNV as identified by bulk WGS analysis (metastasis vs blood). **E** GO Biological Process enrichment analysis of Louvain cluster 2 marker genes in scRNA-seq data (Fig. 1E, adjusted p < 0.05), tested against a background of genes detected in at least 10 cells. P-values were adjusted using the Benjamini-Hochberg method. Dot size reflects gene counts; color indicates adjusted p-value.

To determine whether this concordance extended to individual cells, we examined segmented copy-number profiles. All candidate CTCs included in this analysis carried recurrent tumor-associated alterations, including gains at chromosome 5p and losses at chromosome 2q (Fig. 2B, left). Additional differences, including those on chromosomes 1 and 9, distinguished subsets of candidates. Single PBMCs and the PBMC minibulk lacked comparable alterations (Fig. 2B, right).

We then assessed whether DNA methylation provided complementary evidence of tumor identity using the pretrained pan-cancer classifier from the crossNN framework.^24^ SCLC was the highest-scoring class for every candidate-CTC profile included in the classification analysis (Fig. 2C; Supplementary Table S2). Scores for the highest-ranked class ranged from 0.12 to 0.24, compared with 0.02–0.04 for the second-ranked class. The metastatic EPIC array profile also received SCLC as its highest-scoring class and showed a similar distribution of scores across the reference classes.

Among the PBMC profiles, 42 of 47 received “control (blood)” as the highest-scoring class. Three profiles, A3, A5 and A15, received thymoma as their top prediction. Analysis using published cell type-specific hypomethylation markers identified hypomethylation in NK-cell marker regions in A15, whereas limited coverage prevented clear interpretation of A3 and A5 (Fig. S8).^25^ A19 and A22 received Langerhans cell histiocytosis and “control tissue (inflammatory microenvironment),” respectively. Across PBMC profiles, the five highest-ranked classes for each individual PBMC were associated with blood or immune-rich tissues.

The concordant SCLC ranking of candidate CTCs and metastatic tissue supported a shared tumor-associated methylation profile. The classifier did not contain a dedicated cervical small cell neuroendocrine carcinoma class. The SCLC-associated result was compatible with previously described molecular similarities among small cell neuroendocrine cancers arising at different anatomical sites.^26^

To obtain further genetic evidence, we interrogated five patient-specific nonsynonymous exonic single-nucleotide variants selected from matched tumor–blood WGS using the MutaSeq approach.^18^ Targeted primers were incorporated during transcriptome library preparation to enrich the corresponding cDNA regions. Reads supporting a tumor-specific variant were detected in 10 of the 19 candidates with transcriptomes passing QC: seven had at least one read supporting the KDM6A variant, and three additional candidates had reads supporting the PCYT1B variant (Fig. 2D). Of note, both variants were detected in neither single PBMCs nor bulk blood WGS. The remaining three targeted loci were not covered in any cell. Because this analysis interrogated transcript-derived sequences, lack of coverage did not establish absence of the corresponding genomic variant.

Finally, we investigated the transcriptional identity of the candidates. Of the three clusters shown in Fig. 1E, cluster 0 contained 23 CD45-high and one CD45-intermediate cell, cluster 1 contained 21 CD45-intermediate and one CD45-high cell, and cluster 2 contained all 19 candidate CTCs included in the analysis. Gene Ontology enrichment analysis of cluster-specific marker genes identified terms related to neuronal morphogenesis and axon development in cluster 2, consistent with the tumor’s neuroendocrine phenotype (Fig. 2E; Fig. S10; Supplementary Table S3).

Together, tumor-matched copy-number alterations and mutation-supporting reads, complemented by methylation and transcriptional features, established the tumor origin of the analyzed candidates. We therefore refer to these molecularly characterized cells as CTCs in the subsequent analyses.

### Genetic and epigenetic CTC subpopulations show limited transcriptional separation

Having established CTC identity, we investigated the heterogeneity apparent in the copy-number profiles. Differences in genomic segments, including chromosome 1p, distinguished subsets of CTCs (Fig. 2B), prompting us to examine whether corresponding subdivisions were present in their methylomes.

We used MethSCAn to identify genomic regions whose methylation varied among CTCs.^29^ Dimensionality reduction and Leiden clustering of these variably methylated regions (VMRs) identified two methylation groups (Fig. 3A; Fig. S11). Comparing the groups identified 58 differentially methylated regions (DMRs; adjusted P < 0.05; Supplementary Table S4). Their regional methylation profiles showed differences between the groups despite incomplete coverage in individual cells (Fig. 3B).

**Figure 3.**
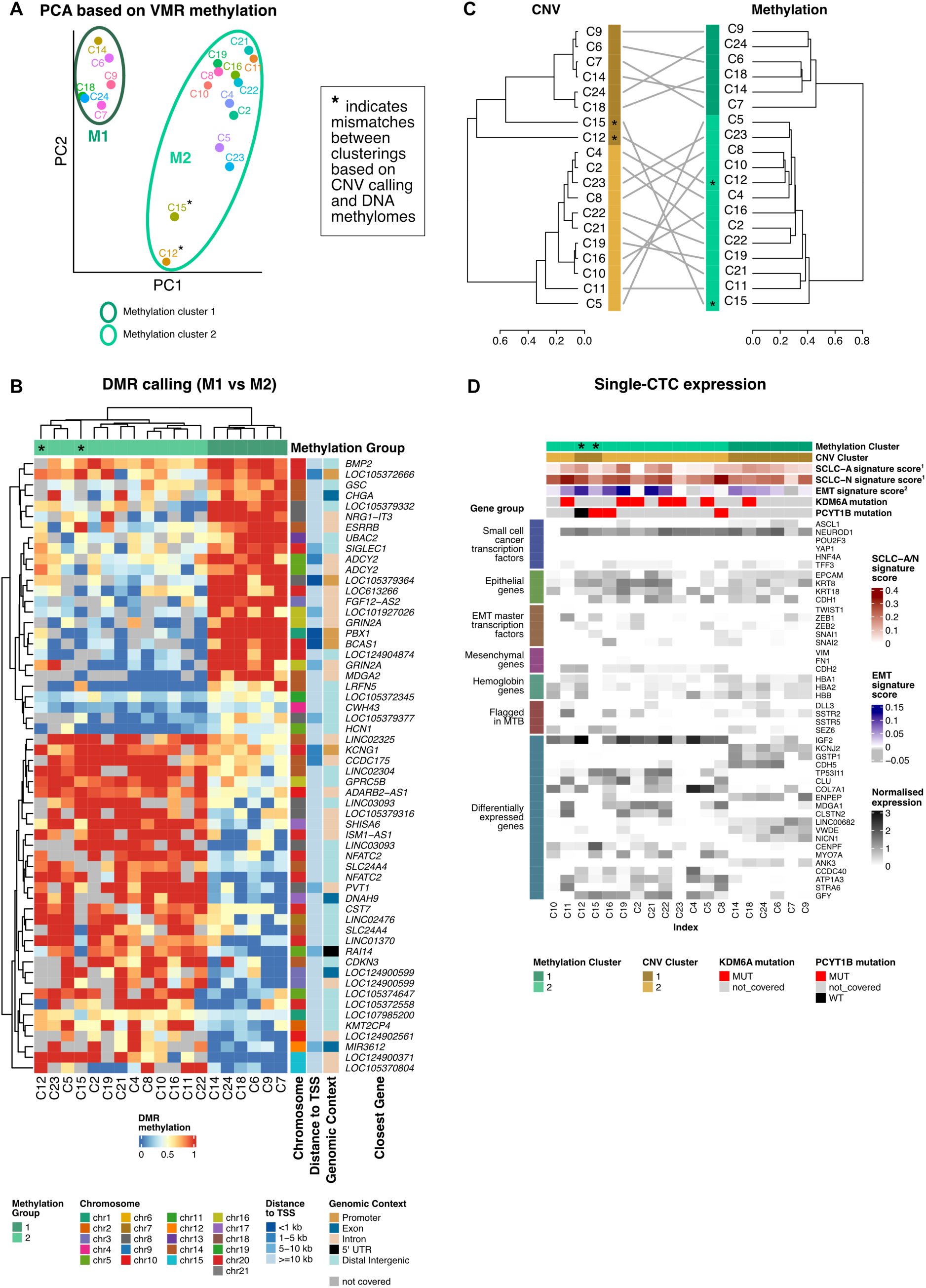
Multi-layer single-cell profiling reveals molecular heterogeneity among CTCs. **A** Principal component analysis of 19 CTCs based on mean shrunken methylation residuals across 66,795 variably methylated regions (VMRs) called with MethSCAn. Missing values were imputed iteratively using a PCA-based approach. Point labels indicate well coordinate. PC, principal component; VMR, variably methylated region. Asterisks mark cells assigned to discordant clusters based on CNVs and DNA methylomes. **B** DNA methylation levels across 58 differentially methylated regions (DMRs) between methylation group 1 (n=6 CTCs) and group 2 (n=13 CTCs) called with MethSCAn. Rows represent DMRs significant at adjusted p-value = 0.05 (multiple testing correction implemented by MethSCAn). Heatmap color indicates mean methylation fraction per DMR. Grey marks regions not covered in a given sample. Row annotations indicate chromosome, distance to nearest transcription start site, and genomic context derived from ChIPseeker annotation against hg38. Gene symbols indicate the closest annotated gene per DMR. Column annotation indicates the cell index and methylation group. TSS, transcription start site; UTR, untranslated region; Chr, chromosome. **C** Comparison of CTC clustering based on copy number variation (CNV) and DNA methylation. Each dendrogram represents hierarchical clustering of 19 CTCs on a single data layer. CNV clustering used 1-Pearson correlation as distance metric on log-transformed segment ratios; methylation clustering used Jaccard distance on VMR methylation values. Both dendrograms used Ward.D2 linkage. Grey lines connect the same cell across the two clusterings. Colored bars indicate cluster membership upon cutting each dendrogram at k = 2. Asterisks mark cells assigned to discordant clusters across the two data layers. **D** Integration of multiple molecular data layers across 19 CTCs. Column annotations show methylation and CNV cluster membership (k = 2 cut, as in C). Signature scores for small cell lung cancer subtypes and epithelial-to-mesenchymal transition (EMT) were calculated using Seurat’s AddModuleScore function based on published gene sets (1 = ref. ^27^, 2 = ref. ^28^). Mutation status at KDM6A and PCYT1B was derived from scIRES-seq transcriptome data; grey indicates insufficient read coverage at the respective locus. The heatmap shows normalized gene expression for a curated panel of marker genes including small cell cancer subtype transcription factors, epithelial markers, EMT master regulators and mesenchymal markers, hemoglobin genes, and genes that were flagged as highly expressed in the molecular tumor board (based on bulk patient data). Using Seurat’s FindMarkers function, differentially expressed genes between cells of methylation cluster 1 vs 2 were identified. The top 20 genes based on absolute log2fold-change are shown. Due to low number of cells, multiple testing correction left no differentially expressed gene with a significant adjusted p-value. Genes are displayed in manually curated order. Columns are ordered by methylation cluster membership. EMT, epithelial-to-mesenchymal transition; MTB, molecular tumor board; MUT, mutant; WT, wild-type.

We next assessed whether the methylation-defined groups corresponded to genetic subpopulations. Hierarchical clustering used log-transformed copy-number segment ratios or methylation profiles of the 1,000 most variable VMRs. When each dendrogram was divided into two clusters, cell assignments agreed for all but two cells (Fig. 3C). Thus, genetic and epigenetic analyses identified largely concordant CTC subpopulations.

To determine whether these subpopulations were accompanied by distinct transcriptional profiles, we examined gene expression alongside CNV group, methylation group and targeted genotype information (Fig. 3D). NEUROD1 transcripts were detected in 18 of 19 CTCs and ASCL1 transcripts in five. These transcription factors are associated with the SCLC-N and SCLC-A transcriptional subtypes, respectively.^27^ Epithelial markers, including EPCAM, KRT8, KRT18 and CDH1, showed variable expression across CTCs. EMT-associated transcription factors, including TWIST1, ZEB1 and SNAI1, showed no consistent expression pattern separating the groups. Detection of the mesenchymal marker CDH2 was restricted to one CTC subpopulation. DLL3, SSTR2, SSTR5 and SEZ6, previously highlighted as highly expressed in the molecular tumor board data, were not detected in every CTC.

Differential expression analysis between the methylation-defined groups identified no genes meeting the adjusted significance threshold after correction for multiple testing. Figure 3D displays the genes with the largest absolute log2 fold changes to illustrate the observed expression differences. Overall, the genetic and epigenetic subdivisions were not accompanied by a clear separation of the measured transcriptional profiles.

Together, these results demonstrate the feasibility of linking phenotypic and molecular measurements to establish tumor identity and characterize heterogeneity in individual patient-derived CTCs. Beyond confirming tumor origin, scIRES-seq revealed substantially concordant genetic and epigenetic subpopulations that were not clearly distinguished by their transcriptomes, illustrating the value of simultaneous multi-omics profiling at single-cell resolution.

## DISCUSSION

This proof-of-concept study establishes scIRES-seq as a framework integrating cell imaging, immunophenotype, copy-number variation, targeted mutation detection, DNA methylation and transcriptomes in individual CTCs. Applied to metastatic cervical small cell neuroendocrine carcinoma, it provided convergent evidence of tumor origin of isolated candidate CTCs and resolved genetic and epigenetic subpopulations. Transcriptomes supported neuroendocrine identity but did not clearly distinguish these subpopulations, illustrating the complementary information obtained through multimodal profiling.

Reliable identification is essential when analyzing rare candidate CTCs among abundant blood-derived events. RTI-SFC enabled direct recognition of debris-like events that fulfilled flow-cytometric gating criteria but lacked intact-cell morphology and did not yield high-quality molecular profiles. Linking images and immunophenotypes to molecular measurements through index sorting therefore provided an additional level of quality assessment. Tumor-matched CNVs established the malignant identity of all analyzed cells, supported by targeted mutations and concordant methylation and transcriptional features. Expanding the dimensions of earlier multimodal CTC studies, scIRES-seq connects such measurements with imaging and immunophenotypic information.^12,13^ Methylation provided complementary evidence where mutation coverage was incomplete, suggesting single-cell methylomes as precise CTC biomarkers, although this case report does not establish methylation alone as a validated diagnostic test.

Most candidate CTCs yielded matched methylome and transcriptome data, supporting the feasibility of the integrated workflow. Its interpretation nevertheless depends on the information accessible to each component. Phenotypic enrichment and viable-singlet selection may exclude other CTC populations. Targeted mutation recovery depends on expression and allelic sampling: three of five selected variants lacked coverage, making their absence from individual cells uninformative.^18^ Sparse methylome coverage primarily limits comparisons at individual CpGs because few positions are consistently covered across cells. Nevertheless, regional analyses resolved methylation heterogeneity, and classification remained concordant across cells with different coverage. These findings support the utility of aggregated methylation information despite incomplete sampling of individual sites.

The SCLC-associated methylation profiles should be interpreted in the context of biological similarities across small cell neuroendocrine carcinomas. Cervical small cell carcinoma is substantially rarer than its pulmonary counterpart: historical US registry data reported age-adjusted incidences of 0.69 and 66.88 cases per million women annually, respectively, during 1992–2010.^30^ This rarity constrains disease-specific molecular studies and makes comparisons with SCLC informative. Shared transcriptional and epigenetic features across small cell neuroendocrine cancers provide a biological basis for such comparisons.^26^ However, cervical and pulmonary small cell carcinomas retain distinct genomic backgrounds, precluding their treatment as biologically interchangeable entities.^20^

In our data, SCLC was the highest-scoring methylation class in both CTCs and metastatic tissue, whereas CESC received low scores. CESC comprises cervical squamous cell carcinoma and endocervical adenocarcinoma and does not represent cervical small cell neuroendocrine carcinoma, which was absent from the reference. The observed classification is therefore compatible with shared neuroendocrine methylation features, without demonstrating that these generally outweigh anatomical origin. Moreover, although crossNN was developed for cross-platform classification of sparse methylomes, its published performance estimates and diagnostic thresholds were not established for single-cell bisulfite profiles.^24^ We accordingly interpret the consistent ranking across CTCs and metastatic tissue as biological concordance rather than a calibrated high-confidence diagnosis.

Beyond tumor identification, scIRES-seq revealed genetic and epigenetic structure within the CTC compartment. The substantial agreement between CNV- and methylation-defined subpopulations suggests an association between genetic background and epigenetic organization. A comparable relationship was observed in colorectal cancer, where methylation patterns tracked genetic lineages while transcriptome groupings did not consistently reproduce them.^31^ Of note, our data do not establish the temporal order or causal relationship between genetic and epigenetic differences. CNVs also provide an incomplete representation of genetic variation, and CNV-associated coverage differences could influence methylation estimates despite the coverage-aware analysis implemented in MethSCAn.^29^ The DMRs characterize groups defined using the same methylation data, hence they do not independently validate those groups or establish regulatory effects on neighboring genes.

The clinical setting further defines the scope of these findings. This study examined one CTC sample from one extensively treated patient and compared it with an earlier metastatic biopsy. The strong molecular concordance supports retained tumor-associated features across these samples, but the absence of primary-tumor data and contemporaneous tissue sampling prevents reconstruction of their evolutionary relationships. Future comparisons of primary tumors, metastases and longitudinal CTC samples could distinguish conserved features from changes accompanying treatment and progression. Blood-based sampling also provides access to disease biology in advanced palliative settings, whose relevance is supported by recent studies of CTC dynamics near the end of life.^32^ Such research requires voluntary participation and procedures adapted to the patient’s condition and goals of care.

The lack of significant transcriptional separation between methylation-defined subpopulations permits several explanations. Small and unequal groups, stochastic transcript detection and multiple-testing correction limit statistical power. Because the groups were identified through methylation, clearer separation in that modality is also partly expected. Biologically, CNVs and methylation differences need not produce strong changes in current gene expression: their effects may be buffered, context-dependent or unrelated to nearby genes. A shared neuroendocrine gene expression program could dominate subtler subgroup differences, while transcriptional variation may follow other axes. Exposure to a common circulatory environment could additionally favor convergent expression programs through adaptation or selection, although this cannot be assessed without appropriate matched tissues and longitudinal comparisons.

Molecular plasticity is another plausible explanation, particularly in light of ASCL1 and NEUROD1 co-expression in a subset of CTCs. Mixed expression of these regulators has been reported in SCLC specimens and cervical small cell neuroendocrine carcinoma.^21,33^ Longitudinal SCLC CTC studies demonstrate treatment-associated changes in transcriptional heterogeneity, while experimental models provide direct evidence of transitions between neuroendocrine programs.^34,35^ Our work in juvenile myelomonocytic leukemia (JMML) provides a complementary example in which postnatal developmental methylation identity coexisted with fetal-like transcriptional programs, with experimental evidence supporting reprogramming.^36^ These observations illustrate how transcriptional activity can diverge from methylation-based identity. In the present study, however, co-expression is compatible with either a stable mixed phenotype or a transitional state. Longitudinal profiling and functional experiments would be required to distinguish plastic adjustment from selection of pre-existing states.

Together, these findings support integrated single-cell profiling as an approach to studying tumor identity and molecular heterogeneity in the circulation. Connecting molecular states to genetic backgrounds could help investigate disease progression, treatment responses and heterogeneous therapeutic targets. The clinical potential of CTC-based target assessment is illustrated by an association between CTC DLL3 protein expression and tarlatamab response in a prospective SCLC study.^37^ Translating scIRES-seq observations into clinical applications will require broader cohorts, longitudinal comparisons and validation linking molecular measurements to protein expression, functional behavior and clinical outcomes.

## METHODS

### Patient samples & ethics approval

The study was conducted according to the Declaration of Helsinki after written informed consent was obtained from the patient. All experiments with human samples were approved by the local Institutional Review Board of Heidelberg University. All information is subject to the provisions of the local data protection regulations. The patient was enrolled in the NCT-MASTER (Molecularly Aided Stratification for Tumor Eradication Research) program, which is an ongoing prospective observational multicentric study (<u>NCT05852522</u>) established by the German Cancer Consortium (DKTK). Within this framework, tumor material and peripheral blood were collected and processed for EPIC methylation array (tumor only) and WGS. All methods related to patient consent, sampling procedure, sample processing, and acquisition, processing and analysis of these data follow standardized workflows and are published in ref.^22^. In addition, a peripheral blood sample for single-cell analysis was collected and processed as described below. Given the patient’s advanced terminal disease, study procedures were limited to the minimum required for the translational analysis and were performed after written informed consent without interfering with symptom-directed clinical care.

### Sample processing

Peripheral blood (18 mL) for single-cell sorting was collected through standard venipuncture into EDTA-K3 tubes (Sarstedt) and kept at room temperature. Processing began within 45 minutes of collection. The PBMC fraction was enriched by gradient centrifugation using equal volumes of Histopaque-1077 (Sigma-Aldrich) and blood, as described in ref. ^38^. The PBMC-enriched fraction was washed with 10 mL ice-cold MACS buffer (DPBS, 0.5 % bovine serum albumin, 2 mM EDTA) and pelleted by centrifugation at 400 x g for 10 min at 4 °C (hereafter “standard centrifugation”). The pellet was resuspended in 2 mL ACK Lysing buffer (Lonza) and was incubated for 5 min at 20 °C for erythrocyte lysis. Lysis was stopped by addition of 10 mL cold MACS buffer, followed by standard centrifugation and resuspension in MACS buffer. CD45+ cells were depleted by magnetic-activated cell sorting (MACS) using LD columns and anti-human CD45 microbeads (both Miltenyi Biotec) according to the manufacturer’s instructions. Both the CD45-depleted flowthrough fraction and the CD45-enriched bead-eluate fraction were collected, and cell counts were determined before immunofluorescence staining.

### Immunofluorescence staining

Cells were stained with a 5-color antibody panel: CD45 PE/Cy7 (Biolegend, #304016), EpCAM PE/Dazzle 594 (Biolegend #324231), CD235a APC, CD44 BV570 (Biolegend, #103037), Vimentin PerCP/Cy5.5 (Biolegend, #677813), diluted 1:50 - 1:100 in MACS buffer. CD45-enriched and CD45-depleted fractions (2 × 10⁶ cells or all available cells, respectively) were incubated with the antibody mix for 30 min at 4 °C in the dark, washed, and resuspended in MACS buffer. Unstained, single-stained (compensation beads), and live/dead (heat-killed, 70 °C, 5 min) controls were included for spectral compensation. DAPI was used to assess viability.

### Real-time imaging spectral flow cytometry

High-speed spectral cell sorting was performed using a BD FACSDiscover S8 Cell Sorter (BD Biosciences / Waters Biosciences, San Jose, CA, USA) equipped with 5 excitation lasers (349, 405, 488, 561, and 637 nm) utilizing a 130-µm nozzle at a system pressure of 7 psi and a drop drive frequency of 15 kHz. High-speed fluorescence imaging was captured simultaneously via integrated BD CellView Image Technology utilizing the 488 nm blue excitation line. Voltage adjustment and spectral unmixing matrices were generated using single-stained OneComp eBeads (Invitrogen) and stained patient PBMCs cells. Unstained patient PBMCs were used to estimate autofluorescence, though these may not fully represent CTC autofluorescence. To minimize sample loss, 20,000 events with images were first recorded for the unstained CD45-enriched, full-stained CD45-enriched, and full-stained CD45-depleted fractions; acquisition was then paused to set the gating strategy (detailed gating strategy in Supplementary Data) based on the recorded fluorescence values. Fluorescence measurements were used for sorting. Images and image-derived parameters were recorded for analysis following sequencing. The target plate was kept at 4 °C throughout sorting. Target populations were sorted in single-cell mode with index sorting enabled. The plate was sealed with aluminized foil seal immediately after sorting, briefly centrifuged and stored at -80 °C until library preparation. Total time from blood draw to freezing the sorted cells was 11h.

### MutaSeq primer design

Genotyping with MutaSeq^18^ requires prior knowledge about the target loci. Somatic SNVs and indels were called within the standardized workflows of the MASTER program. We selected 5 loci as targets for genotyping in this study based on the following criteria: 1) non-synonymous somatic SNV, 2) exonic location of the SNV, 3) high variant allele frequency. PrimerDesign (https://github.com/veltenlab/PrimerDesign) was used to design MutaSeq primers with parameters –nested none –cDNA –optim –blast –readlength 75 –prefixInnerLeft TCGTCGGCAGCGTCAGATGTGTATAAGAGACAG –prefixInnerRight GTCTCGTGGGCTCGGAGATGTGTATAAGAGACAG –numprimers 10. MutaSeq primers are listed in Supplementary Table S1B.

### scIRES-seq library preparation

scIRES-seq was implemented as a 384-well workflow linking index-sort immunophenotypes with DNA methylation, transcriptome and targeted genetic information from individual cells. All data layers are linked through their well coordinate. The workflow combined G&T-seq-based separation of genomic DNA and polyadenylated RNA with adapted scBS-seq and Smart-seq2 library preparation and MutaSeq-based enrichment of selected expressed variants.^14,16,39,17,40,18^ In this study, the workflow was coupled to RTI-SFC as described in sections “Immunofluorescence staining” and “Real-time imaging spectral flow cytometry”.

#### Separation of genomic DNA and polyadenylated RNA

Individual cells were index-sorted into 384-well plates containing 3 µL RLT Plus buffer per well, with the plate holder maintained at 4°C. Plates were centrifuged and stored at −80°C until processing.

Polyadenylated RNA was captured using biotinylated oligo-dT₃₀VN coupled to Dynabeads MyOne Streptavidin C1 (Invitrogen). Beads were prepared using alkaline and salt washes, incubated with the biotinylated oligonucleotide for 15 min at room temperature, washed four times with binding and wash buffer, and resuspended in 1x RT buffer containing 1U/µl recombinant RNase inhibitor (Takara). Beads were used within 30 minutes to capture polyadenylated RNA.

Plates containing cell lysates were thawed on ice right before addition of beads. Each cell lysate received 4 µL bead suspension and underwent three 3-min incubations at room temperature with intervening mixing to capture polyadenylated RNA. Beads were magnetically separated, and the gDNA-containing supernatant was transferred to a corresponding collection well. Two successive 8-µL washes with G&T wash buffer were performed to collect the entirety of the gDNA fraction and transferred to their corresponding collection well. Captured polyadenylated RNA was reverse-transcribed on the beads. Genomic DNA was purified using a 0.7x AMPure XP bead ratio and resuspended in 1 µL EB before bisulfite conversion. The Bravo NGS Workstation was used for all the described steps to minimize sample loss and for accurate pipetting.

#### Reverse transcription and targeted cDNA amplification

Bead-bound polyadenylated RNA was reverse-transcribed in 3 µL master mix containing 6.67 U Maxima H Minus reverse transcriptase (Thermo Scientific), 1 U RNase inhibitor, 1× RT buffer, 1 mM dNTPs, 3 µM template-switching oligonucleotide and 7.5% PEG.

Reverse transcription was performed at 42°C for 90 min, followed by inactivation of the reverse transcriptase at 70°C for 15 min. The alternating 50°C/42°C incubations used in the published SmartSeq2 and MutaSeq protocols were omitted.^18,40^

After reverse transcription, full-length cDNA and selected target regions were co-amplified using 3.15 µl of master mix containing 2x KAPA HiFi HotStart ReadyMix (Roche), 0.16 µM ISPCR primer and a 0.03 µM pool of target-specific primers (see Supplementary Table S1B for primer sequences and targets) carrying Nextera-compatible adaptor sequences.

Master mix was added with the Bravo NGS workstation. Amplification comprised 98°C for 3 min, followed by 22 cycles of 98°C for 20 s, 67°C for 60 s and 72°C for 6 min and a final extension at 72°C for 5 min. Amplified cDNA was purified with a 0.83x ratio of AMPure XP beads and eluted in 7 µL EB. AMPureXP Beads were added with the Mantis system, whereas the subsequent supernatant removals and ethanol washes were performed with the C.Wash system and the elution performed on the Bravo workstation. Representative samples and controls were assessed using TapeStation D5000 analysis.

#### Bisulfite conversion and methylome library preparation

Genomic DNA was bisulfite-converted using Zymo conversion reagents in the presence of the AMPure beads retained from gDNA purification. CT Conversion Reagent (Zymo, #D5003-1) was prepared freshly before use, according to the manufacturer’s’ instructions. 6 µl of conversion reagent were added using the Bravo NGS Workstation to 1 µL of eluted gDNA with AMPureXP beads remaining in the well, followed by incubation at 98°C for 8 min and 65°C for 180 min. Following the incubation steps, the conversion reaction was stored overnight at 4°C.

Converted DNA was purified using 1.5 µl of MagBead J magnetic beads (Zymo, #D4100-3-3) per reaction. Converted gDNA was bound to beads by the addition of 20 µl M-Binding buffer (Zymo) and incubated for 5 minutes. Beads as well as the binding buffer were added to the plates with the Bravo NGS Workstation. Bound gDNA was afterwards washed in M-Wash buffer (Zymo) on the C.Wash and desulphonation with M-Desulphonation buffer (Zymo) was performed for 15 min at room temperature, followed by another two M-Wash buffer washes with the C.Wash. After the last wash, beads were left to dry for 5 minutes at 55°C. Converted gDNA was eluted in 9.7 µl first strand synthesis mix, containing 1.03x Blue Buffer, 0.41 mM dNTPs and 0.41 µM PreAmp_Next_8H primer (see Supplementary Table S1B), and transferred to a fresh plate with the Bravo NGS Workstation.

First strand synthesis used the primer PreAmp_Next_8H, comprising a 5′ C3 spacer, a Nextera-compatible adaptor sequence and an eight-base degenerate region containing A, C or T at each position. Following denaturation at 65°C for 3 min and cooling on ice, 15 U Klenow fragment lacking exonuclease activity (Qiagen, #P7010-HC-L) were added with the Mosquito LV to give an initial reaction volume of 10 µL.

Five rounds of priming and extension were performed. Each round included incubation at 4°C for 5 min, a gradual increase to 37°C in 0.2°C increments every 3 s, and extension at 37°C. Extension lasted 30 min in the first four rounds and 90 min in the final round. Between rounds, samples were denatured at 95°C for 45 s, cooled on ice, and supplemented with 0.6 µl fresh master mix containing 1x Blue Buffer, 4 µM PreAmp_Next_8H, 0.4 mM dNTPs and 10 U Klenow enzyme. Residual primers were digested with 10 U exonuclease I (New England Biolabs, #M0293L) for 60 min at 37°C. The Mosquito LV was used for the addition of the fresh master mix and Exonuclease I.

First strand products were purified with 0.72x AMPure XP beads and resuspended in 12 µl second strand synthesis mix containing 0.41 µM of a Nextera-compatible adaptor primer with a six-base random region (Adapter2_Next_6N), 1.02 x Blue Buffer and 0.41 mM dNTPs. Following denaturation at 95°C for 45 s and cooling on ice, 15 U Klenow enzyme were added. Second strand synthesis used the same gradual 4–37°C temperature increase as in the first strand synthesis, followed by extension at 37°C for 90 min. Double-tagged products were purified by PEG-mediated rebinding to the retained AMPure beads, leading to a 0.73x AMPureXP bead ratio. The Mantis system, as well as the Bravo NGS Workstation and C.Wash were used for the clean-ups and elutions, as well as the Mosquito LV for master mix and Klenow additions.

Methylome libraries were amplified in 12.5 µL reactions containing KAPA HiFi HotStart ReadyMix and unique dual-index primers (see Supplementary Table S1A for UDI sequences). Master mix and primers were added with the Bravo NGS Workstation. Amplification comprised 95°C for 2 min, followed by 14 cycles of 94°C for 80 s, 63°C for 30 s and 72°C for 30 s and a final extension at 72°C for 3 min. Indexed libraries were pooled and subjected to two successive 0.75x AMPure XP bead clean-ups before final quality control.

#### Transcriptome library preparation

Amplified cDNA containing the MutaSeq-enriched targets was used for transcriptome library preparation. Tagmentation employed EMBL Tn5 transposase loaded with Nextera-compatible adaptor oligonucleotides by scOpenLab as described by Seufert and colleagues.^41^

Each tagmentation reaction contained 0.4 µL cDNA with a maximum concentration of 1.5 ng/µl, 0.4 µL of the 1:50 Tn5 preparation, 0.4 µL tagmentation buffer containing 40mM MgCl_2 a_nd Tris-HCl pH 7.5 and 0.4 µl dimethylformamide, giving a total volume of 1.6 µL. Reactions were incubated at 55°C for 10 min and tagmentation was terminated by adding 0.4 µL of 0.1% SDS, followed by incubation on ice for at least 5 min. The Mosquito LV system was used for all pipetting steps.

3 µl master mix containing 1.8x KAPA HiFi HotStart ReadyMix and 10 % DMSO were added to the reaction with the Mantis system as well as 1 µl of 3 µM Unique dual-index primers for library amplification with the Mosquito LV (see Supplementary Table S1A for UDI sequences). Cycling comprised 72°C for 3 min and 98°C for 30 s, followed by 11 cycles of 98°C for 20 s, 63°C for 15 s and 72°C for 30 s and a final extension at 72°C for 3 min.

Indexed libraries were pooled and purified twice using 0.9× AMPure XP beads. The resulting library pool contained both transcriptome-derived fragments and targeted MutaSeq products.

#### Library quality control, indexing and sequencing

Final methylome and transcriptome library pools were quantified using Qubit, and fragment-size distributions were assessed using TapeStation D5000 analysis. A set of 768 unique dual indices was used for library indexing, thus allowing the pooled sequencing of methylome and transcriptome libraries on the same flow cell. Methylome and transcriptome libraries were pooled at a target read ratio of ∼10:1 (methylome:transcriptome) and sequenced on a NovaSeq X Plus (25B flow cell) with paired-end 150 bp reads and dual 10 bp indices (Nextera UDI). Sequencing depth per cell was 1.18 Mio (median raw read pairs) for transcriptome libraries and 0.72 Mio (median raw read pairs) for methylome libraries.

## Data processing

scBS-seq data was processed using nf-core/methylseq v4.2.0 (doi: 10.5281/zenodo.1343417) of the nf-core collection of workflows^42^, utilising reproducible software environments from the Bioconda^43^ and Biocontainers^44^ projects. The pipeline was executed with Nextflow v25.10.0^45^ using the following parameters: --clip_r1 8 --clip_r2 8 –three_prime_clip_r1 8 -- three_prime_clip_r2 8 --local_alignment --non_directional --aligner bismark, and with hg38 as reference genome. Tools used in the workflow included FastQC (https://www.bioinformatics.babraham.ac.uk/projects/fastqc/), MultiQC^46^, TrimGalore! (https://www.bioinformatics.babraham.ac.uk/projects/trim_galore), and Bismark^47^. Methylation calls of the same CpG site from the top and bottom strand were collapsed using methrix v1.20.0^48^. High-quality cells were retained for downstream analyses if they fulfilled the following criteria: 1) minimum 50 % uniquely aligned reads, 2) minimum 100,000 CpG sites covered and 3) maximum 5 % methylation in CHH context.

scRNA-seq data was processed using nf-core/rnaseq v3.15.0 (doi: 10.5281/zenodo.1400710). The pipeline was executed with Nextflow v25.10.0 using the following parameters: –aligner star_salmon –min_mapped_reads 0, with hg38 as a reference genome and the GRCh38 reference annotation (Ensembl release 103, GTF). Tools used in the workflow included STAR^49^, Salmon^50^, StringTie^51^, RSeQC^52^, Qualimap^53^, FastQC (https://www.bioinformatics.babraham.ac.uk/projects/fastqc/), MultiQC^46^, TrimGalore! (https://www.bioinformatics.babraham.ac.uk/projects/trim_galore) and picard (https://broadinstitute.github.io/picard/).

High-quality cells were retained for downstream analyses if they fulfilled the following criteria: 1) minimum 10,000 raw reads, 2) minimum 1000 detected genes and 3) maximum 10 % mitochondrial gene counts.

Genotypes at MutaSeq loci (coordinates provided in Supplementary Table S1B) were called per cell using bcftools^54^ mpileup and call (v1.20). Pileups were generated at predefined target regions from each cell’s BAM file, using a minimum mapping quality of 20, extended BAQ computation, and increased depth limits to avoid coverage capping (-d/-L 10,000,000). Allelic depth (AD) and read depth (DP) were annotated at the FORMAT level. Variants were called using the multiallelic-caller model (bcftools call -m). Per-cell VCFs were sorted, indexed, and merged into a single multi-sample VCF using bcftools merge. vcfR^55^ v1.15 was used for further data handling.

RTI-SFC data was processed with the FACSChorus software v6.3.0. Images were extracted from .cvw files with FlowJo v11. Fluorescence measurements from .fcs files were evaluated in R v4.4.3 using the packages flowCore^56^ v2.18.0, flowWorkspace v4.18.1 (doi:10.18129/B9.bioc.flowWorkspace) and ggcyto^57^ v1.34.0.

### Data analysis

#### Transcriptome analyses

Seurat^58^ v5.3.0 was used for single-cell transcriptome analyses. Mitochondrial and ribosomal gene counts were determined with Add_Mito_Ribo from the scCustomize v3.2.4 package (https://github.com/samuel-marsh/scCustomize). Gene expression data were normalized using log-normalization (scale factor 10,000) as implemented in Seurat. The top 2,000 variable features were identified using the variance-stabilizing transformation (vst) method, and data were subsequently scaled. Principal component analysis (PCA) was performed on the scaled data, and the number of informative principal components was assessed using an elbow plot. A shared nearest-neighbor graph was constructed using the first 20 principal components (k = 15 nearest neighbors). Cells were clustered using the Louvain algorithm (resolution = 0.5) as implemented in Seurat’s FindClusters function, based on the shared nearest-neighbor graph. Normalized gene expression levels, genotype status and Louvain clusters were visualized on the PCA embedding.

Cluster marker genes were identified with the Seurat FindAllMarkers function, with genes restricted to upregulated markers. Only genes expressed in at least 25% of the cells in the cluster and with a minimum log-fold change of 0.25 were considered. Differential expression was assessed using the Wilcoxon rank-sum test. For gene ontology term enrichment analysis (Biological Processes), markers with an adjusted p-value < 0.05 were used with the enrichGO function of the clusterProfiler^59^ package v4.14.6 and org.Hs.eg.db annotation. Benjamini-Hochberg correction was performed, and the background universe was restricted to genes detected in at least 10 cells across the dataset.

An EMT score was calculated with Seurat’s AddModuleScore function using the gene set provided in Table S1A of ref. ^28^. Similarly, module scores for SCLC-A and SCLC-N subtypes were calculated based on gene lists described in ref. ^27^.

#### CNV analysis and clustering

Copy number was assessed with copykit^60^ v0.1.3 largely following the user guide (https://navinlabcode.github.io/CopyKit-UserGuide/). .bam files of scBS-seq data of QC-passing single cells and of bulk tumor and bulk blood WGS were used as input. Variable binning was run with 500 kb resolution, hg38 as reference, removal of the Y chromosome, the CBS method and min_bincount set to 10. The consensus of all PBMCs and of all CTCs was calculated with the median of the “ratios” assay using the calcConsensus function.

CNV tracks were plotted using the single-cell ratio consensus for PBMCs or CTCs respectively, and on the ratio for the bulk tumor or blood sample. For the CNV heatmap, segment ratios called on the full sample cohort were plotted with ComplexHeatmap^61^ v2.20.0. For the dendrogram in Fig. 3C, hierarchical clustering used 1-Pearson correlation as distance metric on log-transformed segment ratios and Ward.D2 linkage.

#### Methylation-based entity prediction with crossNN

CpG sites in the scBS-seq data (for PBMCs and CTCs) or in the EPIC array data (for the metastasis) overlapping the 450K methylation array probes were identified with the corresponding manifest file (HM450.hg38.manifest.tsv, downloaded from https://zwdzwd.github.io/InfiniumAnnotation, ref. ^62^. Methylation beta values >0.6 were encoded as 1, betas <0.6 encoded as -1 and missing values as 0, as described for the crossNN framework^24^. Entity prediction for each sample was performed with the pan-cancer model (https://gitlab.com/euskirchen-lab/crossNN/-/blob/master/models/pancan_devel_v5i_NN.pkl) using the code provided in ref. ^24^. For visualization, only entities receiving prediction rank 1-5 in any sample are shown. As the original publication only determined high-confidence thresholds for bulk data types but not scBS-seq data, we only report raw prediction scores.

#### Differential methylation analysis

scBS-seq data was analyzed with MethSCAn^29^ following the software tutorial. In brief, the functions methscan prepare, methscan filter and methscan smooth were run with the default settings and using quality filtering thresholds as described above. Variably methylated regions (VMRs) were detected with methscan scan and the average methylation in each VMR was calculated with methscan matrix. Missing values in the VMR mean shrunken residuals matrix were iteratively imputed with the package irlba v2.3.5.1 (doi:10.32614/CRAN.package.irlba) based on principal component analysis using 6 principal components given the small number of cells. Leiden clustering of the principal components revealed two clusters between which differentially methylated regions (DMRs) were called with methscan diff. DMRs were filtered for significance based on the adjusted p-value < 0.05. Average DMR methylation was calculated with methscan matrix. Genomic context and nearest gene of each DMR were annotated with ChIPseeker^63^ v1.42.1 using TxDb.Hsapiens.UCSC.hg38.knownGene (doi:10.18129/B9.bioc.TxDb.Hsapiens.UCSC.hg38.knownGene) as reference.

#### Cell type specific methylation analysis

A methylation atlas with 250 marker regions specifically unmethylated in one cell type was used for this analysis (file “Markers.U250.hg38.tsv” from https://github.com/nloyfer/UXM_deconv, ref. ^25^). Methylated and total read counts per cell and marker region were determined with the methscan matrix function. Read counts were summed up per cell for marker regions belonging to the same cell type, and were used to calculate average beta values. Marker regions with low coverage (< 30 covered CpGs) were highlighted visually.

### AI Use

Generative AI (Claude Sonnet 4.6, Anthropic; GPT-6 Astra, OpenAI) was used during the preparation of this manuscript to assist with code generation and debugging for data analysis scripts, as an entry point for literature searches, to synthesize narrative text from author-provided bullet points, and to support revision of manuscript text for clarity and concision. All AI-assisted output was reviewed, verified, and edited by the authors, who take full responsibility for the content, analyses, and conclusions of this work.

## Code and Data availability

Gene counts and beta values per single cell will be made available in the GEO database upon publication of the manuscript in a peer-reviewed journal.

The code used for data analysis and visualization is provided upon request.

## Acknowledgements

We are grateful to the patient for the willingness to contribute to research while receiving palliative care. We also thank the palMA palliative care unit and the patient advisory board at University Hospital Mannheim for supporting our research. We thank NCT Heidelberg for providing the molecular tumor board data used in this study. We thank all members of the Lipka, Scholl and Fröhling groups for critical discussions and valuable feedback. We thank the scOpenLab, the Flow Cytometry Core Facility, the NGS Core Facility, and the Omics IT and Data Management Core Facility at the German Cancer Research Center (DKFZ), for providing excellent services. We thank Cihan Erkut for support on data management. This work was in part funded by Health+Life Science Alliance to M. Hartmann.

## Author Contributions

**Conceptualization:** Mark Hartmann, Daniel B. Lipka, Tobias Boch, Jan-Philipp Mallm. **Formal analysis:** Lea Kubitz, Valentin Maurer, and Daniel Hübschmann. **Funding acquisition:** Mark Hartmann, Daniel B. Lipka, Tobias Boch, Stefan Fröhling, Jan-Philipp Mallm. **Investigation:** Lea Kubitz, Katharina Bauer, Maximilian Schönung, Panna Lajer, and Mark Hartmann. **Methodology:** Lea Kubitz, Katharina Bauer, Diana Ordoñez-Rueda, Maximilian Schönung, Valentin Maurer, Panna Lajer, Stephen J. Clark, Steffen Schmitt, and Mark Hartmann. **Project administration:** Mark Hartmann. **Resources:** Wolf-Karsten Hofmann, Steffen Schmitt, Jan-Philipp Mallm, Stefan Fröhling, Tobias Boch, Daniel B. Lipka, Mark Hartmann, Daniel Hübschmann, and Maria-Veronica Teleanu. **Supervision:** Mark Hartmann, Daniel B. Lipka, Stefan Fröhling. **Visualization:** Lea Kubitz and Mark Hartmann. **Writing - original draft:** Mark Hartmann, Daniel Lipka, Lea Kubitz, Katharina Bauer, and Tobias Boch. **Writing - review & editing:** All authors.

## Conflicts of interest

All authors declare no conflict of interest.

## Notes

### Competing Interest Statement

The authors have declared no competing interest.

