## Supplementary Figures for "scIRES-seq reveals multimodal molecular heterogeneity in CTCs from small cell cervical cancer"

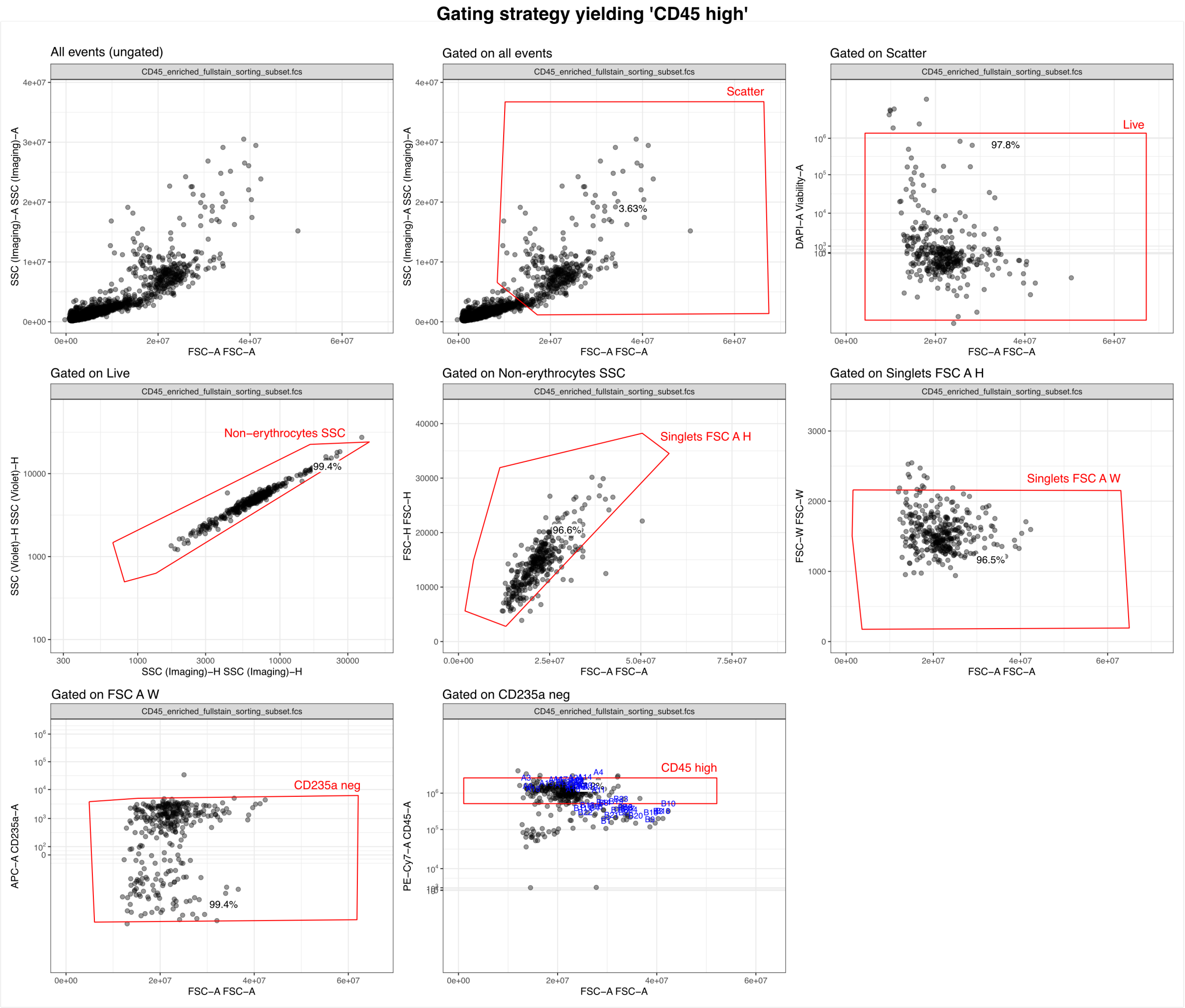


**Figure S1. Gating strategy for sorted CD45-high PBMCs.** Red polygons enclose the target population and were set based on marker expression and scatter properties. PBMCs were sorted from the CD45-enriched fraction after MACS. For visualization, .fcs files were randomly downsampled to 1000 raw events including the sorted events. Blue dots indicate sorted events and the respective well coordinate.

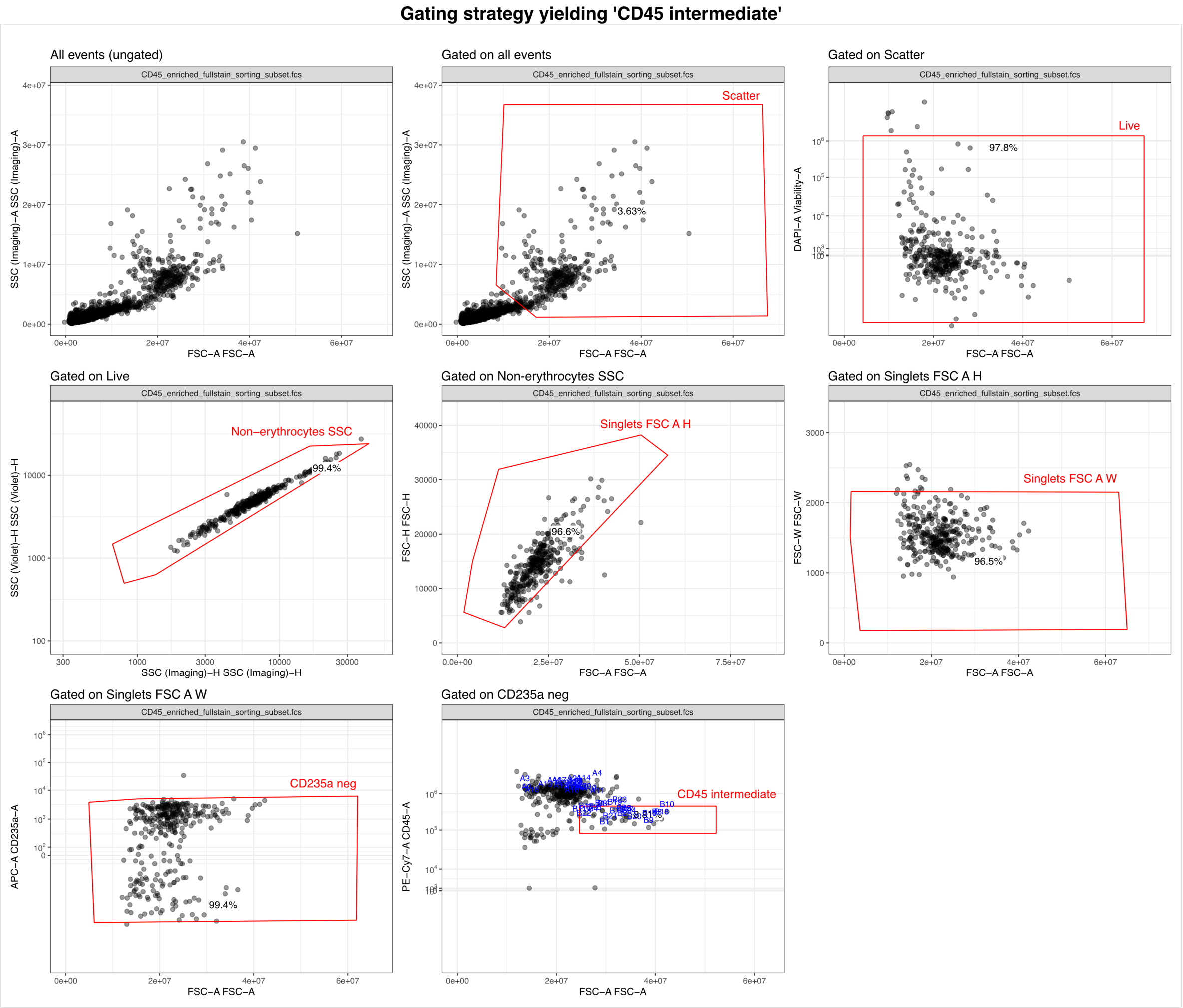


**Figure S2. Gating strategy for sorted CD45-intermediate PBMCs.** Red polygons enclose the target population and were set based on marker expression and scatter properties. PBMCs were sorted from the CD45-enriched fraction after MACS. For visualization, .fcs files were randomly downsampled to 1000 raw events including the sorted events. Blue dots indicate sorted events and the respective well coordinate.

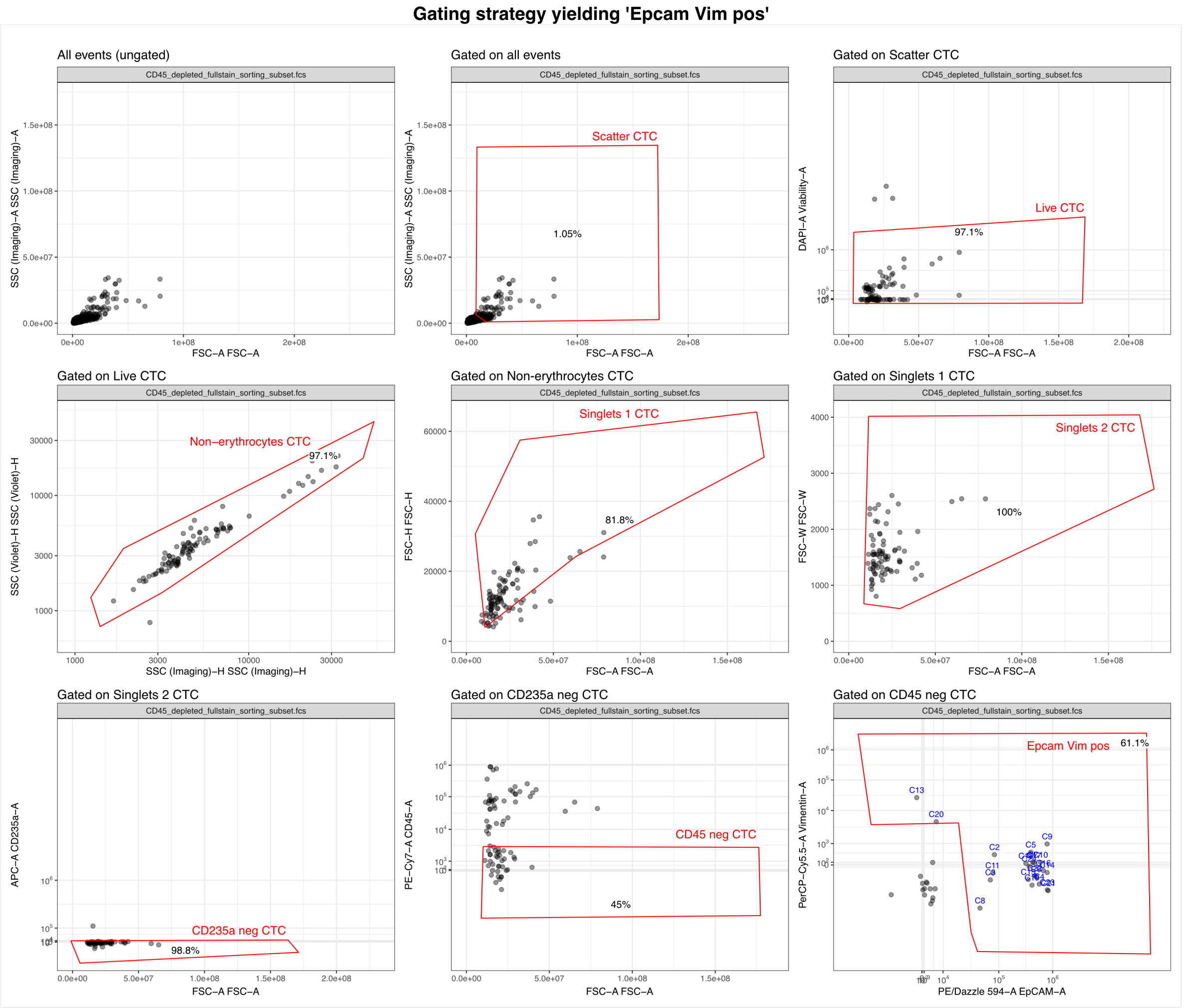


**Figure S3. Gating strategy for sorted candidate CTCs.** Red polygons enclose the target population and were set based on marker expression and scatter properties. Candidate CTCs were sorted from the CD45-depleted fraction after MACS. For visualization, .fcs files were randomly downsampled to 1000 raw events including the sorted events. Blue dots indicate sorted events and the respective well coordinate.

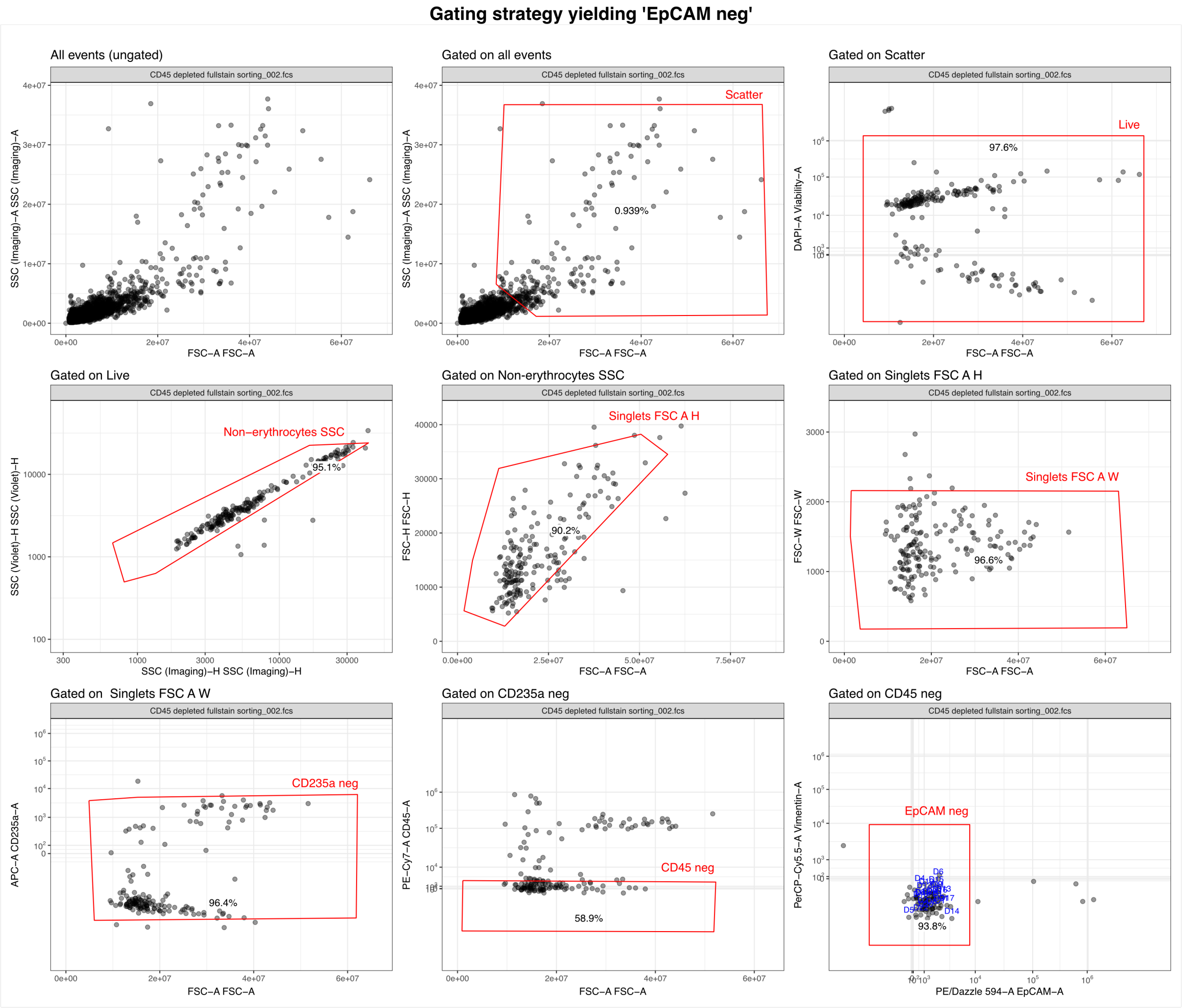


**Figure S4. Gating strategy for sorted debris.** Red polygons enclose the target population and were set based on marker expression and scatter properties. Debris events were sorted from the CD45-depleted fraction after MACS. For visualization, .fcs files were randomly downsampled to 1000 raw events including the sorted events. Blue dots indicate sorted events and the respective well coordinate.

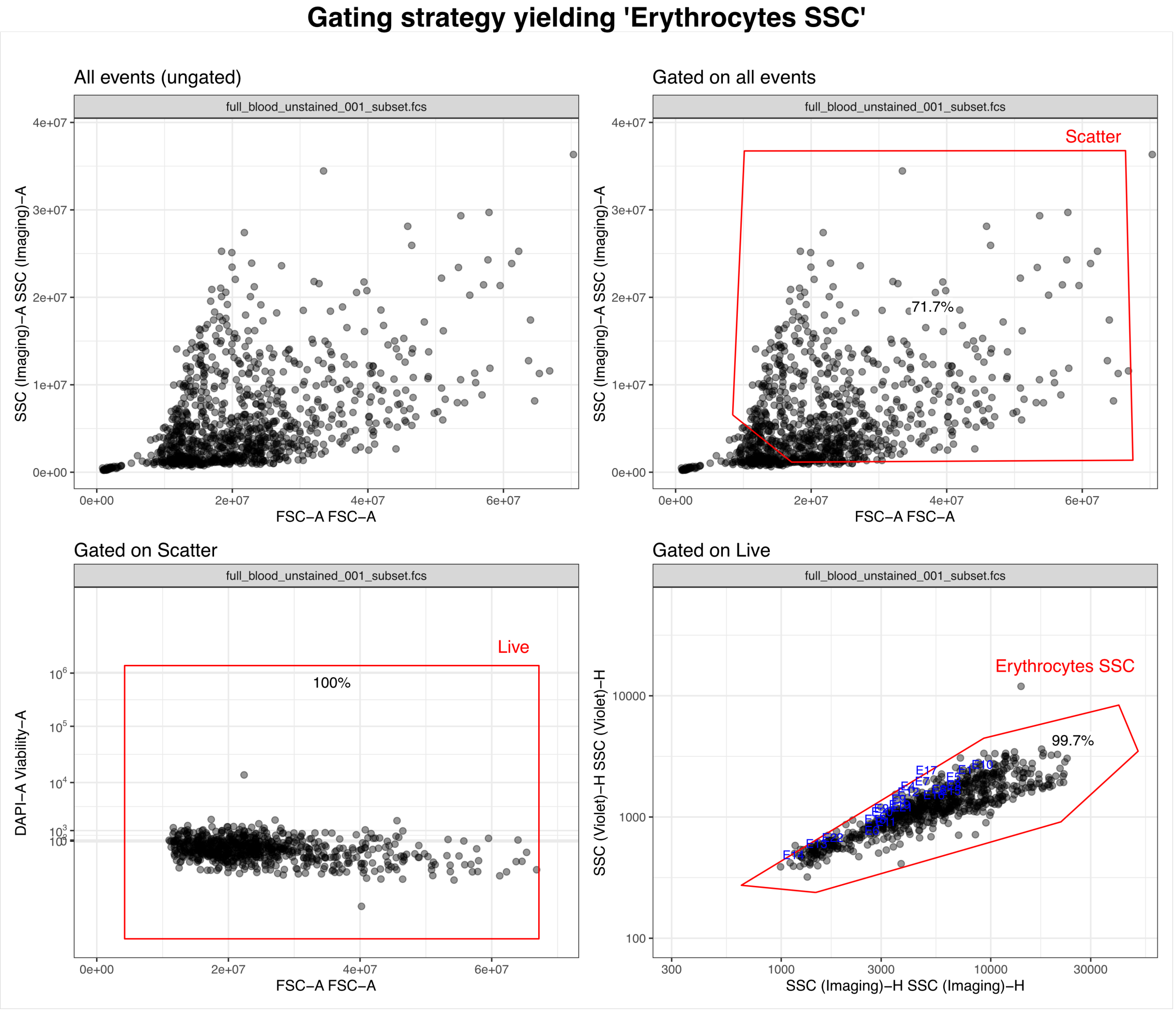


**Figure S5. Gating strategy for sorted erythrocytes.** Red polygons enclose the target population and were set based on scatter properties alone following the protocol described in ref. [64]. Erythrocytes were sorted from unstained whole blood. For visualization, .fcs files were randomly downsampled to 1000 raw events including the sorted events. Blue dots indicate sorted events and the respective well coordinate. FSC-A, forward scatter-area; SSC-H, side scatter-height; PBMC, peripheral blood mononuclear cell.

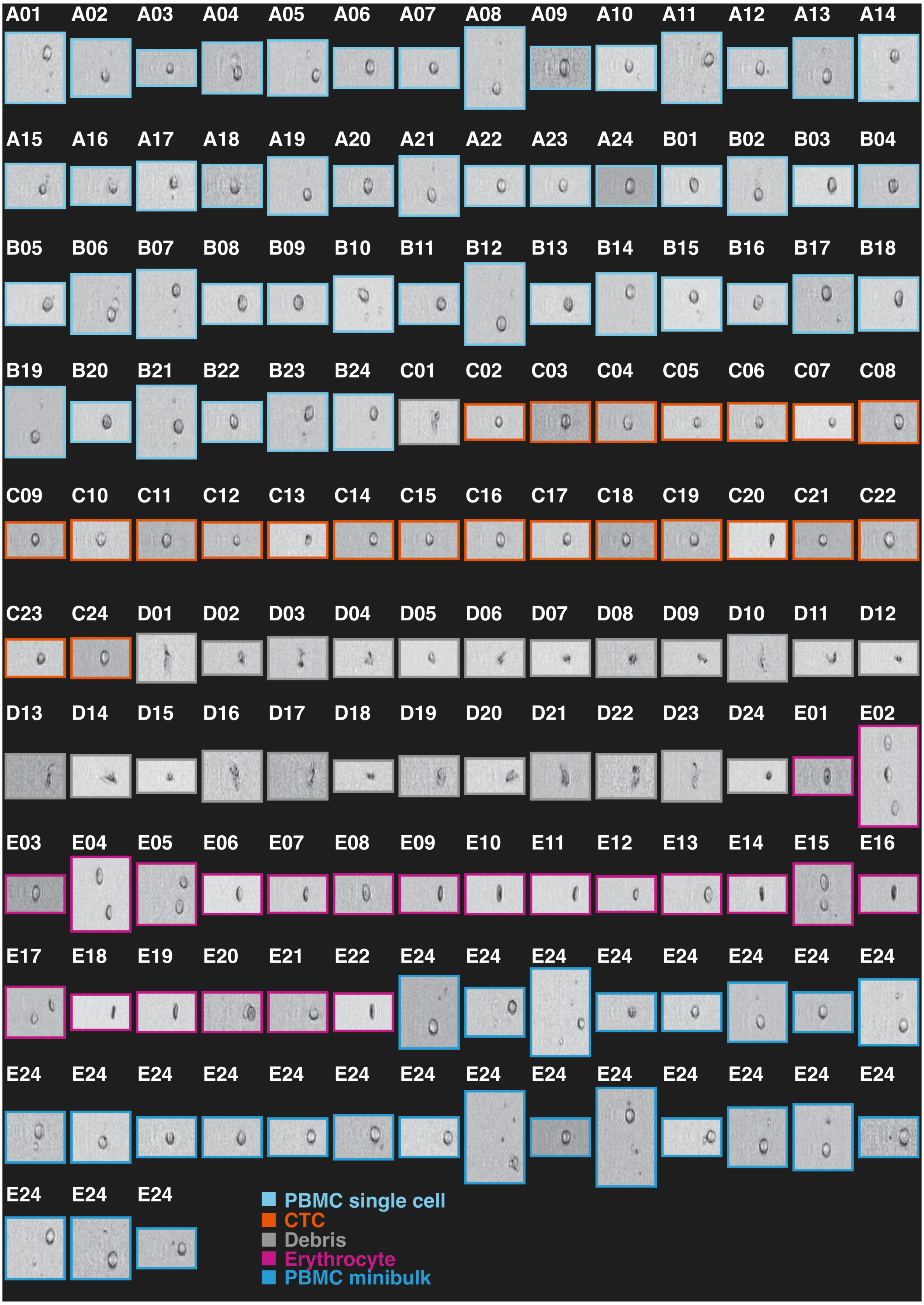


**Figure S6. Images of all sorted events.** Data were acquired with the Lightloss channel of the FACSDiscover S8 instrument. Widths of the images are consistent between all events. The CellView technology does not provide scale bar information. Colored frames indicate sorting population.

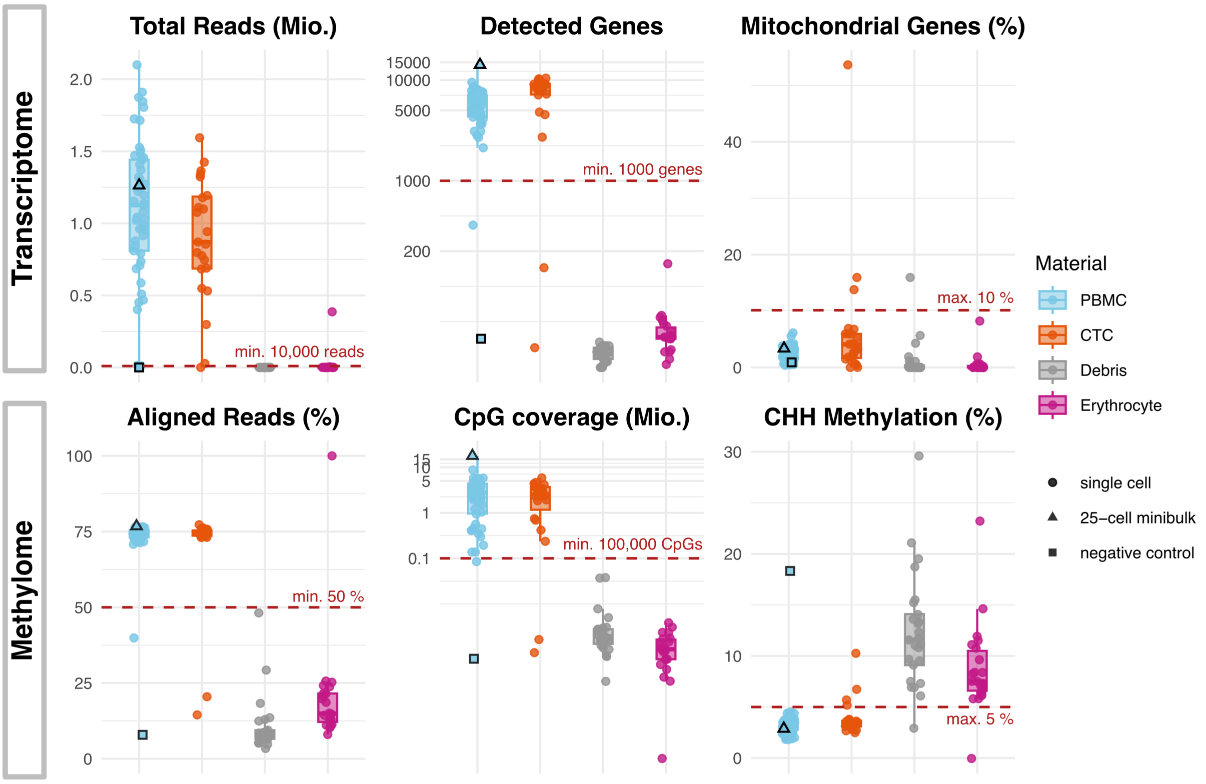


**Figure S7. Quality control metrics of all sorted events.** Red dashed lines indicate cutoffs. Top row transcriptome metrics, bottom row methylome metrics. Positive control (25-cell minibulk) and negative control (no cell) are highlighted with black triangle and square respectively. Boxes show the IQR, center lines the median and whiskers extend to 1.5 × IQR.

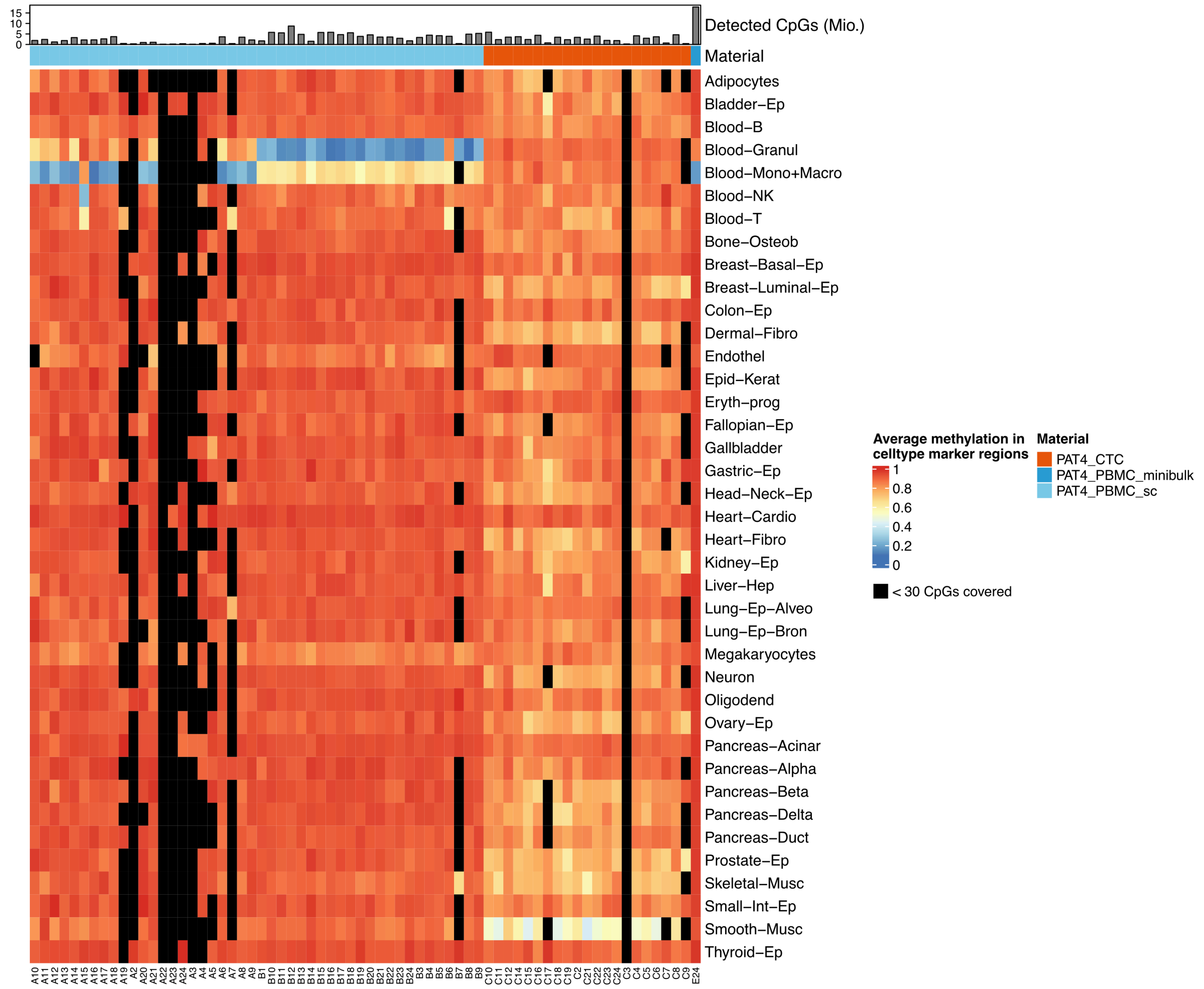


**Figure S8. Average methylation in cell type specific marker regions across samples.** Color indicates mean methylation fraction. Per cell type, 250 marker regions were obtained from ref. [25]. Rows and columns were hierarchically clustered using Euclidean distance and complete linkage. Black rectangles indicate marker regions in which less than 30 CpG sites were covered the respective sample. Column annotations indicate the sample material. Column names refer to well coordinates. PBMCs with coordinates A and E were sorted from the CD45-high gate, while cells with coordinates B were sorted from the CD45-intermediate gate.

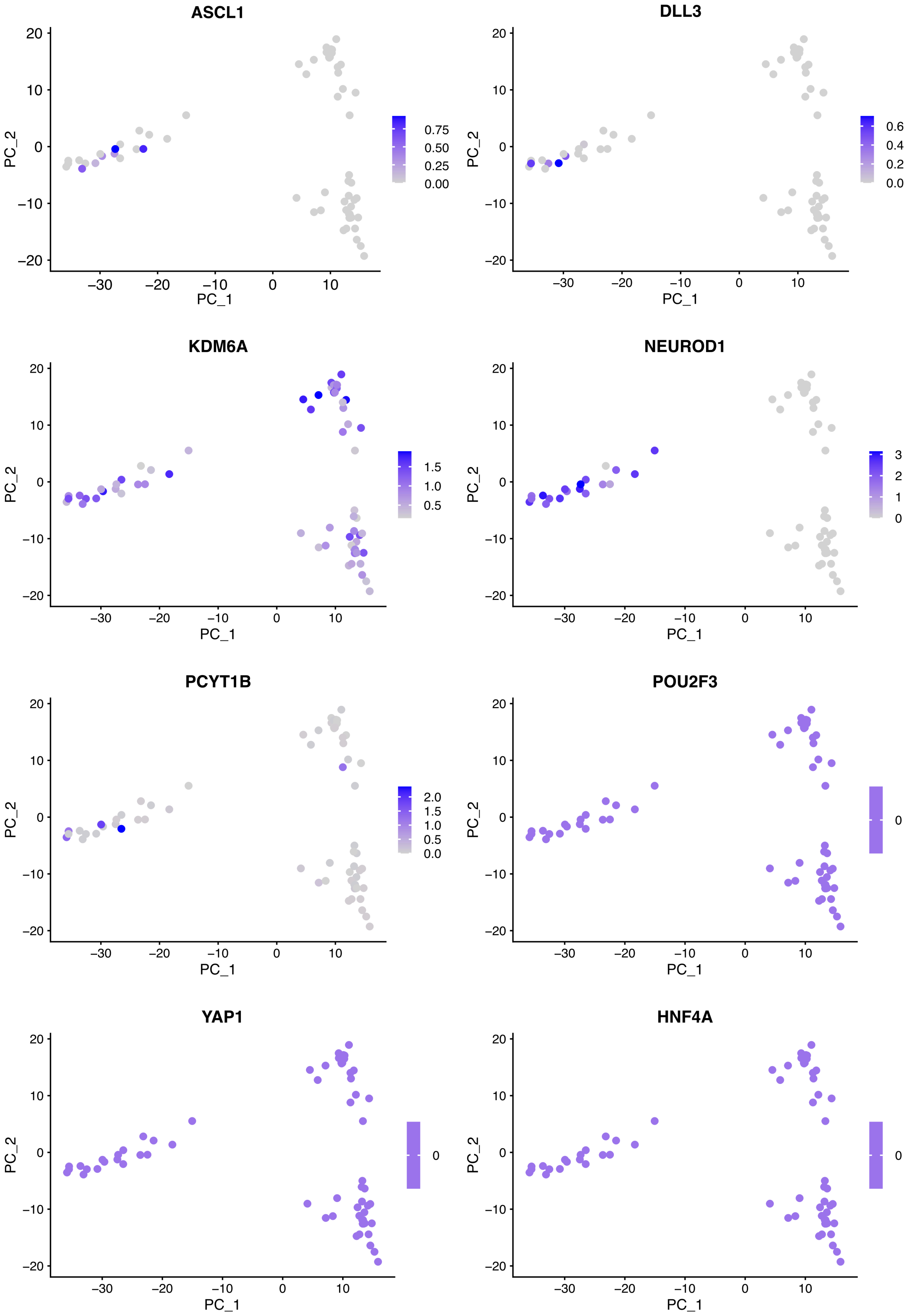


**Figure S9. Principal component analysis of single-cell transcriptomes.** Color indicates the log-normalized expression of the gene listed in each plot title.

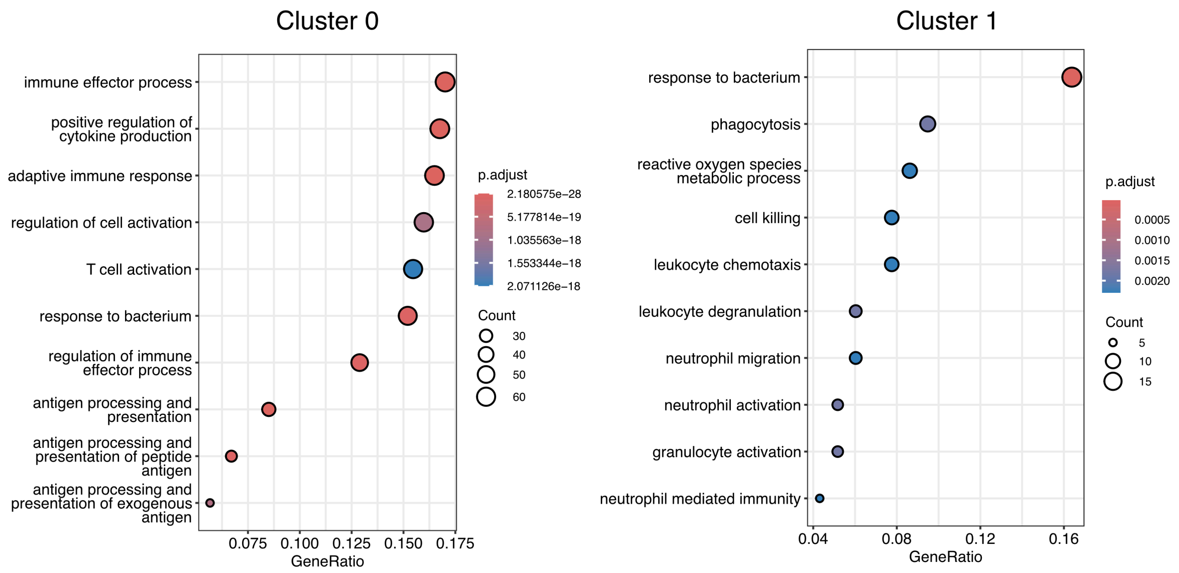


**Figure S10. GO Biological Process enrichment analysis of Seurat clusters 0 and 1 marker genes.** Adjusted $p < 0.05$ (clusters from Fig. 1E), tested against a background of genes detected in at least 10 cells. P-values were adjusted using the Benjamini-Hochberg method. Dot size reflects gene counts; color indicates adjusted p-value.

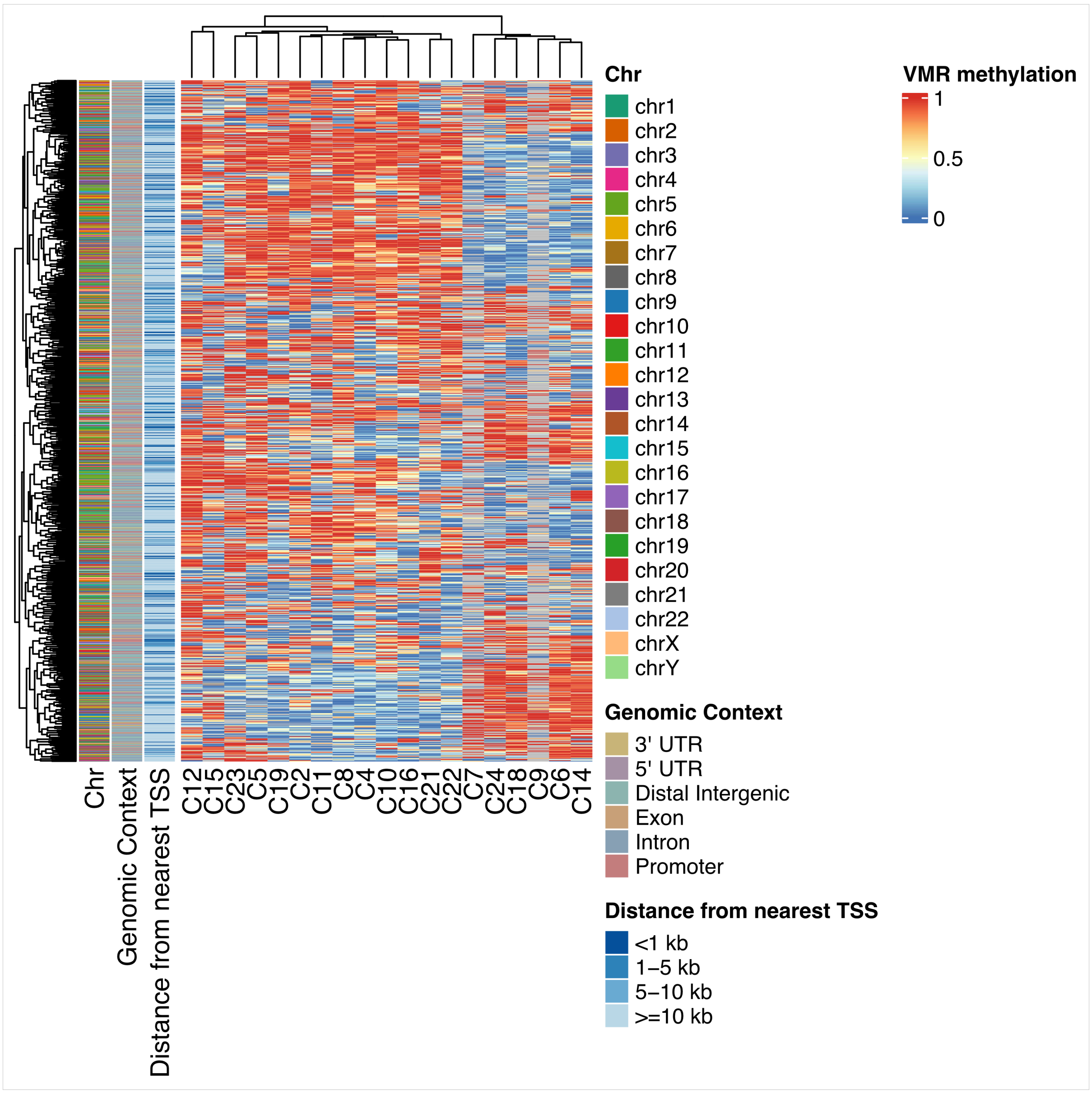


**Figure S11. Average methylation in variably methylated regions (VMRs) called on all CTCs with MethSCAn.** Rows are regions, columns are cells. All VMRs (n = 66,795) were filtered for coverage in minimum 10 cells. 1000 VMRs with the highest variance of the filtered VMR subset are plotted. Color indicates average methylation fraction and grey shows regions without coverage in the respective cell. Row annotations show chromosomal location of each VMR, its genomic context and its distance to the nearest transcription start site. Rows and columns were hierarchically clustered using Euclidean distance and complete linkage. TSS, transcription start site; UTR, untranslated region; Chr, chromosome.
